# Oral antigen exposure under co-stimulation blockade generates Treg cells to establish immune tolerance despite prior sensitization

**DOI:** 10.1101/2025.04.02.646742

**Authors:** Masaya Arai, Ryoji Kawakami, Yamami Nakamura, Yoko Naito, Daisuke Motooka, Atsushi Sugimoto, Tomiko Kimoto, Naganari Ohkura, Norihisa Mikami, Shimon Sakaguchi

**Affiliations:** Department of Experimental Immunology, Immunology Frontier Research Center, The University of Osaka, 3-1, Yamadaoka, 565-0871, Suita, Osaka, Japan; Department of Experimental Immunology, Institute for Life and Medical Sciences, Kyoto University, Sakyo-ku, 606-8507 Kyoto, Japan; Genome Information Research Center, Research Institute for Microbial Diseases, The University of Osaka, 3-1, Yamadaoka, 565-0871, Suita, Osaka, Japan; Department of Frontier Research in Tumor Immunology, Graduate School of Medicine, The University of Osaka, 3-1, Yamadaoka, 565-0871, Suita, Osaka, Japan

## Abstract

Antigen-specific oral tolerance is effective in preventing harmful immune responses in antigen-non-sensitized animals but difficult to be induced in those already antigen-primed. Here, we show in mice that feeding of antigen-containing diet generates peripherally derived regulatory T (pTreg) cells that exhibit a tissue-adapted effector phenotype. The antigen feeding also enables the generated pTreg cells to acquire Treg-specific epigenomic changes in Treg signature genes including *Foxp3*, hence the stability of Treg-specific function. Cessation of antigen feeding, however, results in decline in oral tolerance with diminution of pTreg cells. Transcriptomic analysis has revealed that the induced pTreg cells predominantly express CD101. CD101^+^ Treg cells with a similar phenotype and epigenetic alterations can also be generated *in vitro* from antigen-primed naïve CD4^+^ T cells by blockade of CD28-mediated co-stimulation during TGF-β-dependent Treg induction. Furthermore, *in vivo* CD28 signal blockade by CTLA-4-immunoglobulin fusion protein (CTLA4-Ig) before antigen feeding can establish oral tolerance in the mice precedingly antigen-sensitized by a non-oral route. The blockade facilitates differentiation of antigen-specific conventional T cells into CD101^+^ pTreg cells. Thus, continuous oral antigen administration combined with CD28 co-stimulation blockade generates CD101^+^ antigen-specific functionally stable pTreg cells, which can establish long-term systemic antigen-specific immune tolerance even in antigen-pre-sensitized animals.

## Introduction

Oral tolerance is antigen-specific systemic immunological unresponsiveness induced by intestinal exposure to antigens derived from ingested food or commensal microbes (Chase, 1946; Cerovic et al., 2024). It is effective in preventing autoimmune diseases (e.g., arthritis, diabetes, and multiple sclerosis) and allergies (e.g., food allergy) in animal models (Rezende and Weiner, 2022). In humans, early exposure to peanut protein is reportedly capable of preventing peanut allergy in children (Du Toit et al., 2015, 2024). However, it has not been significantly successful in humans to induce oral tolerance for treating ongoing immunological diseases such as chronic allergy and autoimmune disease (Faria and Weiner, 2005). It is therefore a long outstanding issue to determine how harmful immune responses that have been already triggered and chronically progressing can be halted and cured by inducing antigen-specific oral tolerance (Mowat, 2018).

Naturally occurring CD4^+^ regulatory T (nTreg) cells expressing the transcription factor Foxp3, including thymus-derived Treg (tTreg) cells and peripherally derived Treg (pTreg) cells, are indispensable for the induction and maintenance of immunological tolerance, including oral tolerance, and instrumental in controlling immunological diseases (Sakaguchi et al., 2020; Mucida et al., 2005; Josefowicz et al., 2012; Tanoue et al., 2016; Hong et al., 2022). It remains obscure, however, how antigen-specific pTreg cells are generated and how they establish and maintain oral tolerance (Cerovic et al., 2024). It has been suggested that, following antigen transportation through the intestinal epithelial cells, antigen-presenting cells (APCs), conventional dendritic cells (cDCs) in particular, play a pivotal role in the generation of antigen cognate pTreg cells in mesenteric lymph nodes (mLNs) and lymphoid tissues in the intestine (Mazzini et al., 2014; Hadis et al., 2011; Esterházy et al., 2016). Intestinal cDCs can efficiently convert TGF-β into its active form, and vitamin A into retinoic acid (RA); TGF-β and RA synergistically induce intestinal pTreg cells upon antigen stimulation (Mucida et al., 2007; Sun et al., 2007; Coombes et al., 2007). These pTreg-inducing cDCs appear to be immature DCs with low expression of the costimulatory molecules CD80 and CD86 (Morelli and Thomson, 2007; Benson et al., 2007; Lutz et al., 2021). However, the roles of CD28-mediated co-stimulation for pTreg generation and oral tolerance have not been well characterized. Addressing the issue, we have recently shown that *in vitro* antigen stimulation of CD4^+^ conventional T (Tconv) (i.e., CD4^+^CD25^−^Foxp3^-^) cells in the presence of TGF-β and IL-2 but without CD28 signal is able to efficiently induce Foxp3 Treg cells equipped with Treg-specific epigenetic changes, hence Treg cell-lineage stability, and that such functionally stable *in vitro* induced Treg (iTreg) cells can be generated not only from naïve (i.e., CD4^+^CD25^−^Foxp3^-^ CD62L^hi^CD44^lo^) but also from effector/memory (i.e., CD4^+^CD25^−^Foxp3^-^CD62L^lo^CD44^hi^) Tconv cells (Mikami et al., 2020). The question then arises whether such *in vitro* generation of iTreg cells functionally similar to nTreg cells shares a common developmental basis with *in vivo* pTreg generation, especially in the context of oral tolerance.

Here, we have attempted to determine whether a functionally and phenotypically distinct type of pTreg cells are generated upon antigen exposure via the oral route, how they survive upon antigen stimulation and maintain their function to sustain oral tolerance, and how they can be exploited to establish robust oral tolerance not only in antigen-nonsensitized naïve mice but also in those that have been already antigen primed at other sites of the body.

## Results

### pTreg generation and maintenance by continuous antigen feeding and failure in establishing oral tolerance by prior antigen sensitization

To analyze the development of pTreg cells by distinguishing them from tTreg cells, we used OVA-specific DO11.10 (DO) TCR transgenic mice that were made Rag2-deficient and expressing Foxp3-eGFP reporter gene. DO Rag2KO Foxp3-eGFP mice thus produced are hereafter abbreviated as DORe mice. Foxp3^+^ (i.e., GFP^+^) cells were absent in the thymus, very few in the spleen and lymph nodes including mLNs in untreated DORe mice (see also below). To determine the effects of OVA feeding on anti-OVA T-cell responses in DORe mice, we kept feeding them for 4 weeks with the diet containing egg white powder (EWP) as 10% of the protein content or with control normal chow (NC), and assessed ear swelling after 4 times of tape stripping (TS) and papain/OVA treatment on the right ear and control TS/papain treatment on the left ear (**Fig. 1A**). Compared with NC-fed mice, which showed in the right ears severe ear swelling and histologically evident inflammation in the epidermis and dermis with abundant cellular infiltration, EWP-fed mice exhibited much lower degrees of ear swelling and histologically evident inflammation (**Fig. 1B** and **1C**). They had much smaller numbers of DO CD4^+^ T cells (**Fig. S1A**) and much lower ratios of IFN-γ, IL-4, or IL-17A-producing DO CD4^+^ T cells in the draining lymph nodes (dLNs) (**Fig. S1B**). They developed pTreg cells in mLNs, dLNs, and the right ear in much higher ratios compared with NC-fed mice (**Fig. 1D**). Treg cells were found to be present in the spleen and also in the control left ear as well at high ratios, although in small numbers (**Fig. S1C** and **1D**), with no significant ear swelling or inflammation in EWP-fed mice (**Fig. S1D** and **S1E**). In addition, Diphtheria toxin (DTx) treatment of EWP-fed DO11.10 Rag2KO Foxp3-DTR-GFP (DORF) mice expressing the diphtheria toxin receptor (DTR) specifically in pTreg cells (**Fig. 1E**) depleted them in the OVA-challenged ear and dLNs. It evoked severe ear swelling and inflammation (**Fig. 1F** and **1G**) with significant increases of inflammatory cytokine-producing T cells especially in the ear and dLNs (**Fig. S1F** and **S1G**).

**Fig. 1.**
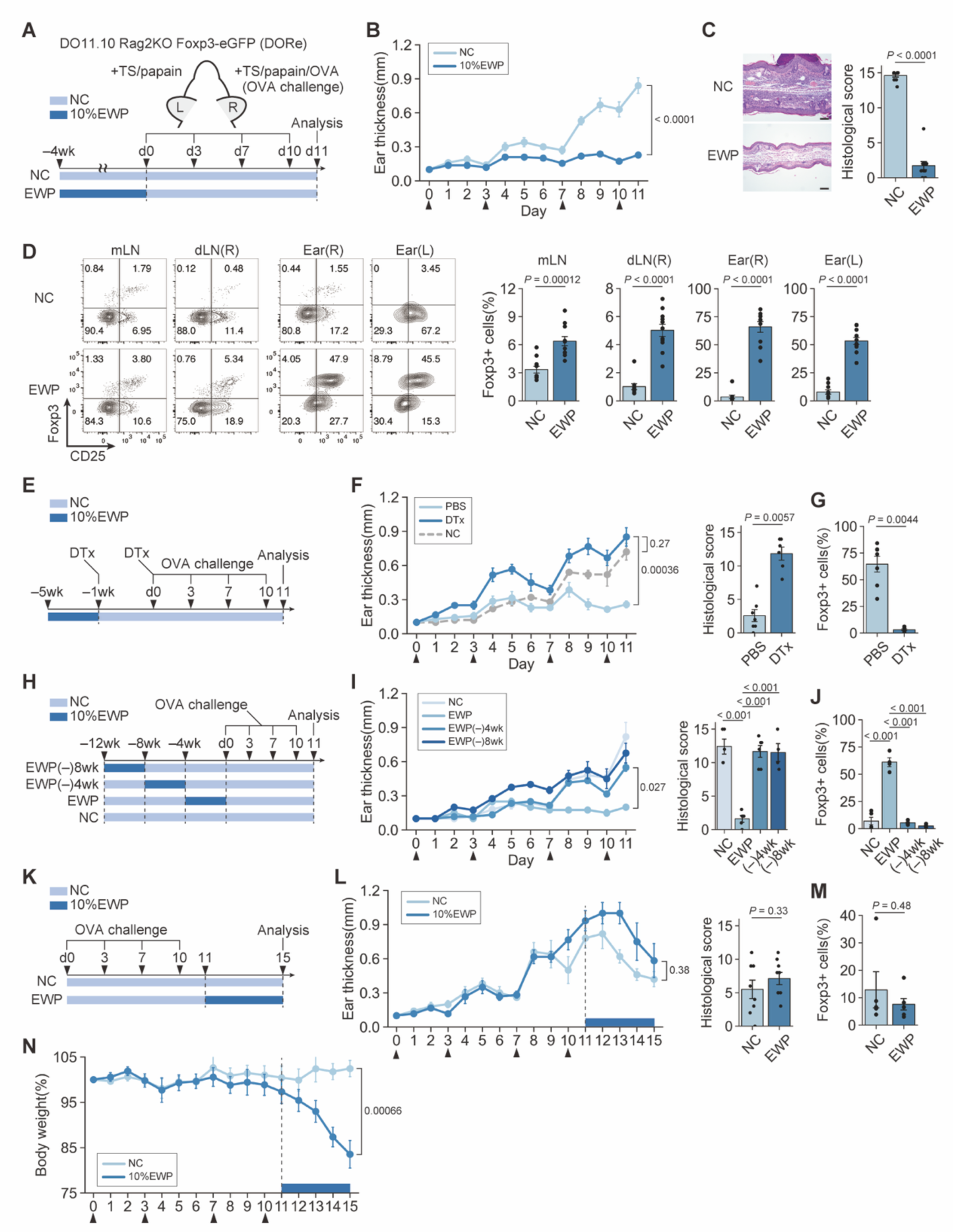
pTreg generation and maintenance by antigen feeding in establishing oral tolerance. (**A**) Antigen feeding to DORe mice and assessment of ear swelling upon antigen challenge. (**B**) Changes in right ear thickness in DORe mice treated as shown in (A). Arrowheads indicate OVA challenges. (**C**) Histology of right ear and histological scores in NC- or EWP-fed DORe mice. Bars in histology indicate 100 µm. (**D**) CD25 and Foxp3 expression by DO CD4^+^ T cells in mLNs, dLNs, and both ears. Barplots indicate frequencies of Foxp3^+^ cells (pTreg cells) (*n* = 10 and 11). (**E**-**G**) Treg cell depletion in EWP-fed DORF mice. Mice were treated twice with DTx followed by OVA challenge (E). Ear thickness and histological scores (F). Frequencies of pTreg cells in the right ear (G) (*n* = 5-7). (**H-J**) Cessation of EWP feeding for 4 weeks (designated EWP(-)4wk) or 8 weeks (EWP(-)8wk) before OVA challenge (H). Ear thickness and histological scores (I). Frequencies of pTreg cells in right ear (J) (*n* = 4-6). (**K**-**N**) Effects of skin OVA sensitization and subsequent OVA feeding on ear swelling upon OVA challenge (K). Ear thickness and histological scores (L). Frequencies of pTreg cells in right ear (M) (*n* = 5 and 6). Changes in body weight of DORe mice (N). Vertical bars indicate mean ± SEM. Statistical significance was assessed by unpaired *t* test or Welch’s *t* test (B, C, D, F, G, L, M, N), and Tukey-Kramer test (I, J).

To determine whether induction of oral tolerance would require continuous antigen exposure, we ceased EWP feeding for 4 or 8 weeks before the OVA challenge to the ear (**Fig. 1H**). Despite prior EWP feeding for 4 weeks, the feeding-suspended DORe mice showed ear inflammation as severe as control NC-fed mice (**Fig. 1I**), reduced ratios of pTreg cells, and increased cytokine-producing T cells in the ear tissue and dLNs when compared to DORe mice that had been kept EWP-fed until the ear testing (**Fig. 1J, S1H**, and **S1I**).

Next, to examine whether antigen-specific oral tolerance could be induced in precedingly antigen-primed mice, we first sensitized DORe mice by TS/papain/OVA treatment and then fed the mice with EWP or NC (**Fig. 1K**). Both groups developed similar degrees of ear swelling and inflammation (**Fig. 1L**). The frequencies of pTreg cells and Tconv cells producing inflammatory cytokines were comparable in mLNs, dLNs, and the ear tissues in both groups (**Fig. 1M, S1J**, and **S1K**). Moreover, the EWP-fed group showed significant body weight loss (**Fig. 1N**), indicating elicitation of systemic inflammation by oral antigen feeding in antigen-sensitized mice.

Collectively, continuous feeding of the diet containing protein antigen establishes antigen-specific oral tolerance by generating pTreg cells from antigen-specific Tconv cells. The generated pTreg cells migrate through the whole body to various tissues. However, prior antigen sensitization at other sites or discontinuity of antigen feeding hampers oral tolerance induction and may even evoke an aberrant immune response upon antigen feeding.

### Antigen-dependent development of pTreg cells with Treg type epigenome in oral tolerance

To examine how the quality and quantity of antigens contained in the diet would affect the development and immunological characteristics of pTreg cells in oral tolerance, we fed DORe mice with the chow containing various concentrations of EWP, antigen-free AIN-93G chow containing only casein as protein (Reeves et al., 1993), or NC, starting from 2 weeks of age before weaning. EWP feeding increased the frequency of pTreg cells in mLNs and the small intestine lamina propria (SI-LP) in an antigen dose-dependent manner, even if antigen-containing chow was taken *ad libitum* (**Fig. 2A** and **2B**). High doses of EWP increased CD25 expression by the pTreg cells induced in SI-LP (**Fig. 2A** and **2C**), indicating their enhanced activation status (Miyara et al., 2009). The EWP-induced pTreg cells were also maintained as CD44^hi^CD62L^lo^ effector/memory type T cells in SI-LP (**Fig. S2A** and **S2B**). These pTregs exhibited Treg-specific epigenetic modifications, including DNA hypomethylation at Treg signature gene loci, such as *Foxp3*-conserved non-coding sequence 2 (CNS2) region and other regions in *Ikzf2*, *Ikzf4*, and *Ctla4* loci (Ohkura et al., 2012) (**Fig. 2D**). By chromatin immunoprecipitation sequencing (ChIP-seq) of the genomic regions associated with histone H3 Lys27 acetylation (H3K27ac) as an indicator of active enhancer status, differential peak analysis revealed that Treg-specific enhancer regions were activated in pTreg cells from EWP-fed DORe mice, whereas Tconv-specific enhancer regions were not, as in nTreg cells in WT mice (Kitagawa et al., 2017; Kawakami et al., 2021; Dikiy et al., 2021) (**Fig. 2E**). The pTreg cells thus generated had activated Treg-specific super-enhancers at the *Foxp3* gene locus and other Treg signature gene loci including *Ikzf2*, *Ikzf4*, *Ctla4*, and *Il2ra* as observed in WT-nTreg cells (Kitagawa et al., 2017) (**Fig. 2F** and **2G**). ATAC-seq further confirmed chromatin remodeling also in these loci (**Fig. 2F** and **2G**).

**Fig. 2.**
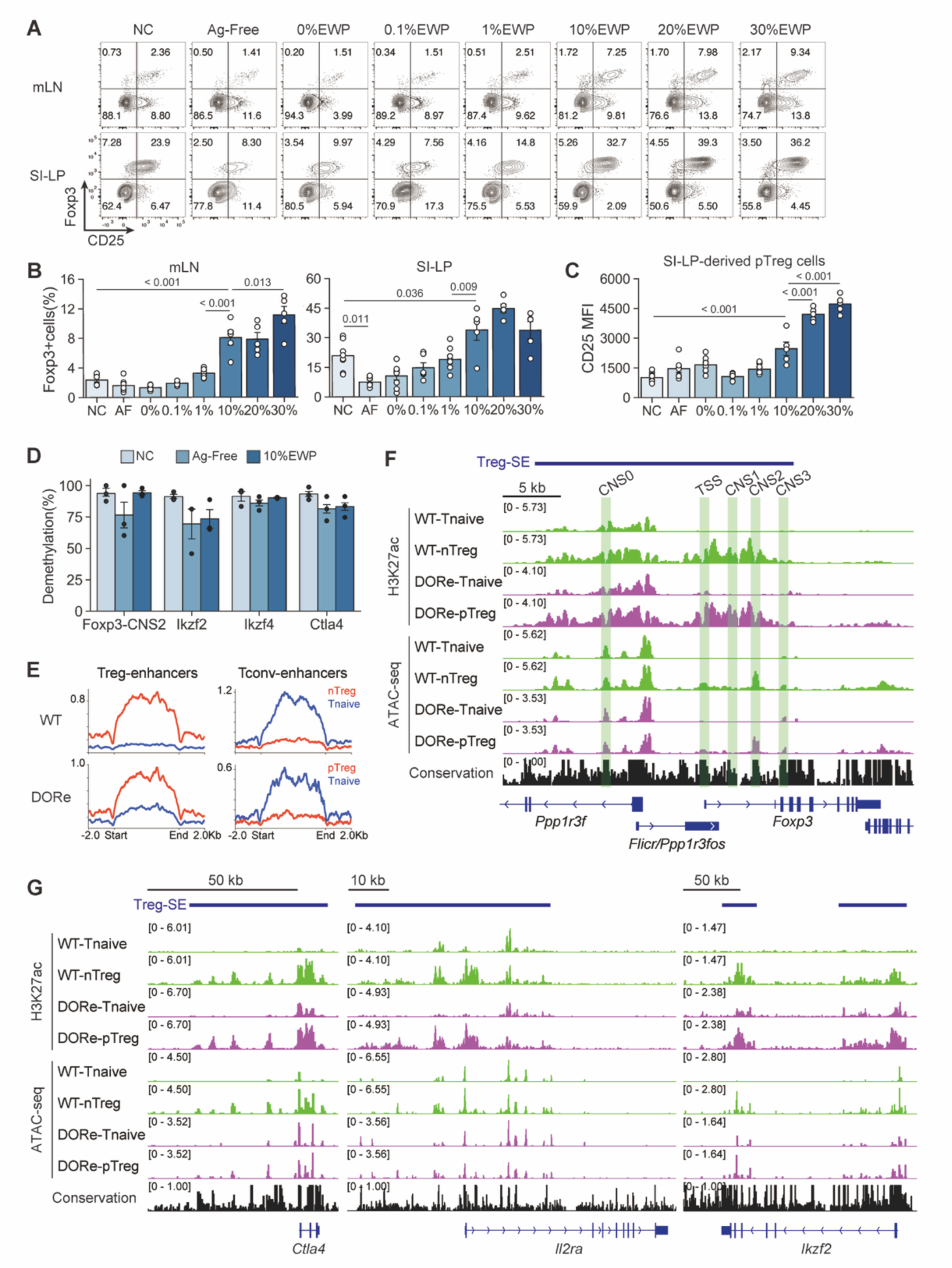
Antigen-dependent development of pTreg cells with Treg type epigenome in oral tolerance. (**A**) Representative CD25 and Foxp3 expression by DO CD4^+^ T cells in mLNs and SI-LP from DORe mice maintained by designated diet for 6 weeks. (**B, C**) Frequencies of Foxp3^+^ cells among DO CD4^+^ T cells (B) and CD25 expression level of Foxp3^+^ cells (C) in mLNs and SI-LP in the mouse groups as shown in (A) (*n* = 5-8). (**D**) Degrees of demethylation as percentage of demetylated CpG residues at Treg-specific demethylation regions in *Foxp3*-CNS2, *Ikzf2*, *Ikzf4*, and *Ctla4* loci of pTreg cells in the mice as shown in **A**. (**E**) H3K27ac ChIP-seq signals at global Treg-specific or Tconv-specific enhancer regions in WT or DORe-derived naïve Tconv or Treg cells. (**F**, **G**) H3K27ac ChIP-seq and ATAC-seq peak call at the *Foxp3* gene locus (F) and other Treg-specific enhancer loci at *Ctla4*, *Il2ra*, and *Ikzf2* (G) in eFox-derived naïve Tconv and nTreg cells, DORe-derived naïve Tconv and pTreg cells. Vertical bars indicate mean ± SEM. Statistical significance was assessed by Tukey-Kramer test (B, C).

In the experiments in Fig. 2A, we noted that NC-fed mice also possessed detectable numbers of pTreg cells in mLNs and SI-LP in particular, at higher frequencies than antigen-free- or 0% EWP chow-fed mice (**Fig. 2A** and **2B**). To analyze the finding, we compared pTreg development in DORe and WT mice after NC feeding, and found that the former developed Foxp3^+^ cells at higher ratios than the latter in SI-LP, the large intestine lamina propria (LI-LP), and among small intestine intraepithelial (SI-IE) cells, but not in the spleen and LNs (**Fig. S2C** and **S2D**). The Foxp3^+^ cells in NC-fed DORe mice exhibited DNA hypomethylation at Treg-specific demethylation regions in Treg signature genes (**Fig. S2F**), and expressed Helios and Neuropilin-1 (**Fig. S2E**), at comparable levels as WT nTreg cells. The results suggested that an antigen(s) cross-reactive with OVA in NC might activate DO Tconv cells to differentiate into functionally stable pTreg cells, although the number of pTreg cells thus generated was not sufficient to induce OVA-specific oral tolerance (**Fig. 1B** and **1C**). After depletion of these pTreg cells in NC-fed DORF mice by DTx administration **(Fig. S2G** and **S2H)**, EWP feeding efficiently induced pTreg cells, suggesting that the pTreg cells were derived from non-Treg cells upon EWP antigen stimulation, rather than from selective expansion or survival of the pre-existing pTreg cells **(Fig. S2I)**.

Taken together, these results indicate that the diet containing cognate antigen drives Tconv cells to differentiate into activated pTreg cells in the intestine, and enables them to acquire Treg-specific epigenetic changes, hence Treg-lineage stability similar to that of nTreg cells. In addition, the quantity of pTreg cells is dependent on the amount of the antigen contained in the diet; thus, more than a certain amount of the antigen is required for generating pTreg cells quantitatively sufficient to establish oral tolerance.

### Antigen feeding induces tissue-specific effector-type pTreg cells suppressing effector Tconv cell differentiation

We next attempted to characterize the dynamics of the differentiation/activation of pTreg and Tconv cells in EWP-fed DORe mice using single-cell RNA-sequencing (scRNA-seq). Analysis of preprocessed 6,449 cells from four hash-tagged and pooled populations of DO^+^ CD4^+^ T cells (i.e., mLN or SI-LP cells in DORe mice NC- or 10% EWP-fed for 4 weeks) identified 13 clusters by the expression of T-lineage signature genes (**Fig. 3A** and **S3A**). EWP feeding resulted in an increase of Cluster 8 (circulating 2), which was *Myb*^hi^ (indicative of augmented T-cell survival and exhaustion) (Yuan et al., 2010; Dias et al., 2017; Tsui et al., 2022), reductions of Cluster 4 (Teff-mix) and 11 (Th1), and conversion of Treg population from Cluster 5 to Cluster 3 in SI-LP (**Fig. 3B**). Compared to Cluster 5, Cluster 3 pTreg cells having increased after EWP feeding highly expressed activation-related genes (*Tnfrsf9*, *Tnfrsf4*, *Tigit,* and *Tnfrsf18*), Th2-type transcription factor (*Gata3*), and tissue repairing-related molecules (*Areg* and *Penk*) (**Fig. 3C**). Supporting these results, gene ontology analysis between Cluster 3 and 5 SI-LP pTreg cells revealed that Cluster 3 pTreg cells expressed a highly activated and proliferative T cell phenotype compared to Cluster 5 pTreg cells (**Fig. 3D**). To analyze further the kinetics of antigen-specific T cells in SI-LP, we re-clustered Cluster 3, 4, 5, and 11 based on commonalities and newly identified 8 clusters (**Fig. 3E-G** and **S3B**). Together with pseudotime analysis (**Fig. 3G**), the analysis suggested that, among Foxp3^+^ cells (Clusters 0, 1, 4, 6, and 7), Clusters 0 and 7 became activated and differentiated upon EWP stimulation into Cluster 1 and 4, which expressed *Klrg1*, *Areg*, and *Penk*. Among Tconv cells (Clusters 2, 3, and 5), effector T-cell populations with Th1 and Th17 gene signatures appeared to diminish upon EWP feeding (**Fig. 3E**, **3F**, and **S3B**). *Mcl1*, an anti-apoptotic gene, was relatively high in *Sell*^+^ (i.e., CD62L^hi^) Foxp3^+^ cells (Cluster 7); *Bcl2* being high in Th1 cells (Cluster 5) (**Fig. S3C**).

**Fig. 3.**
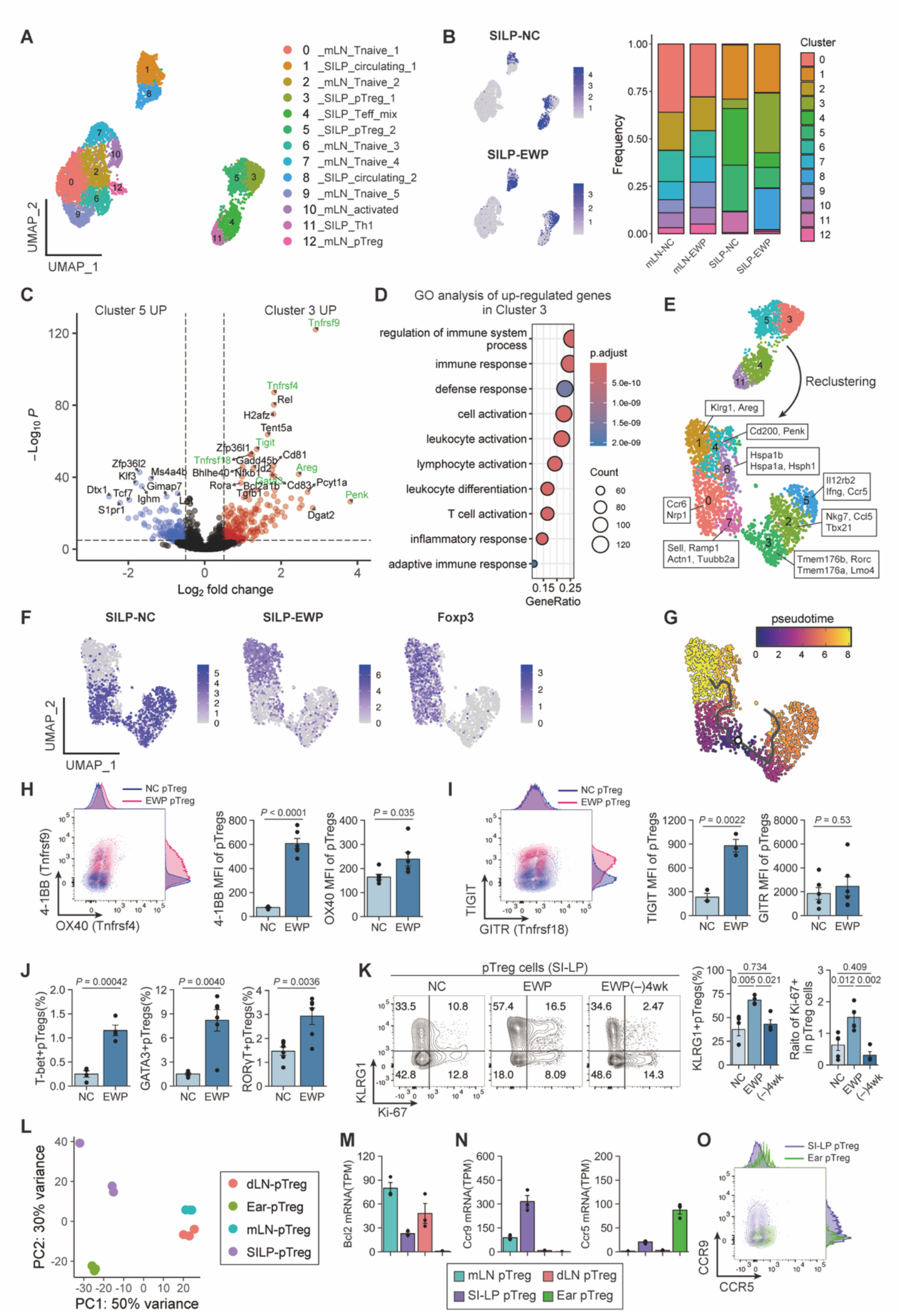
Generation of tissue-specific effector-type pTreg cells by antigen feeding. (**A**) scRNA-seq of DO CD4^+^ T cells in mice fed with NC- or EWP-food for 4 weeks from weaning. (**B**) Hashtag oligonucleotide (HTO) classification of DO CD4^+^ T cells in SI-LP in NC- or EWP-fed mice (left) with proportion of cells in each cluster (right). (**C, D**) Volcano plot of differentially expressed genes (C) and gene ontology analysis (D) between Cluster 3 and 5 pTreg cells. (**E**) UMAP plots of cells re-clustered from Clusters 3, 4, 5, 11 shown in (A), with projection of specifically expressed genes. (**F**) UMAP plots with NC- or EWP-derived HTO and *Foxp3* expression, as shown in E. (**G**) Pseudotime trajectory of re-clustered cells in (F). (**H, I**) Contour plots and MFI of 4-1BB and OX40 (H), TIGIT and GITR (I) staining of pTreg cells in SI-LP of NC- or EWP-fed mice (*n* = 3-6). (**J**) Frequencies of T-bet^+^, GATA3^+^, or RORγT^+^ pTreg cells in NC- or EWP-fed mice (*n* = 4-6 each). (**K**) Ki-67 and KLRG1 expression of SI-LP-derived pTreg cells in DORe mice kept fed with NC or EWP, or ceased to be EWP-fed for 4 weeks EWP (-)4wk as shown in Fig. 1H (*n* = 4-5). (**L**) PCA of transcriptomes of pTreg cells from mLNs, dLNs, SI-LP, and ear tissue (*n* = 3 each). (**M,** and **N**) TPM-normalized *Bcl2* (M), *Ccr9* and *Ccr5* (N) expression by pTreg cells from designated tissues. (**O**) CCR9 and CCR5 expression by SI-LP- and ear tissue-derived pTreg cells. Vertical bars indicate mean ± SEM. Statistical significance was assessed by unpaired *t* test or Welch’s *t* test (H, I, J), and Tukey-Kramer test (K).

We assessed protein expression to validate these findings by comparing EWP- and NC-fed SI-LP pTreg cells (**Fig. 3H-K**). The former showed significantly higher expression of 4-1BB (*Tnfrsf9*), OX40 (*Tnfrsf4*), and TIGIT (*Tigit*), in good correlations with their mRNA expression levels (**Fig. 3H** and **3I**). EWP-fed pTreg cells also contained increased proportions of T-bet^+^, GATA3^+^, or RORγt^+^ pTreg cells compared with NC-fed pTreg cells (**Fig. 3J**). Furthermore, continuous EWP feeding induced Ki-67^+^ and KLRG1^+^ pTreg cells (i.e., highly proliferative and differentiated pTreg cells), which were significantly reduced in ratio after cessation of the feeding for 4 weeks (**Fig. 3K**).

We also conducted bulk mRNA-seq to assess the characteristics of pTreg cells in lymphoid and non-lymphoid tissues in EWP-fed mice. pTreg cells in mLNs and skin dLNs possessed more common features compared to those in SI-LP or the ear (**Fig. 3L**), suggesting that pTreg cells generated in the intestine underwent tissue-specific adaptation and acquired a tissue-specific Treg phenotype and function distinct from those of lymph node pTreg cells, as observed in scRNA-seq (**Fig. S3D**) (Miragaia et al., 2019). For example, SI-LP and ear-tissue pTreg cells (corresponding to Clusters 0, 1, 4, 6, and 7 in Fig. 3E) were distinct from LN pTreg cells in the expression of apoptosis-related molecules (**Fig. S3E**), for example, their lower expression of *Bcl2*, hence being more prone to die by apoptosis after their antigen-specific activation (**Fig. 3M**). Further, among chemokine receptors controlling tissue migration, SI-LP and ear-tissue pTreg cells distinctly expressed CCR9 and CCR5, respectively, in both the mRNA and protein levels (**Fig. 3N, 3O,** and **S3F**).

Collectively, these results demonstrate that EWP feeding generates and sustain pTreg cells that can differentiate into tissue-specific effector-type Treg cells that suppress the differentiation/activation of effector Tconv cells.

### Specific expression of CD101 by intestinal pTreg cells generated by antigen feeding

We next searched for specific markers that could differentiate EWP-induced intestinal pTreg cells from other Treg cells, particularly from tTreg cells, by preparing mixed bone marrow chimeric (BMC) mice in which DORe BM-derived pTreg cells and WT (Thy1.1-eFox) mouse BM-derived nTreg cells developed in the same environment (**Fig. 4A**). While NC or 10% EWP feeding for 4 weeks from immediately after bone marrow transfer equally generated WT nTreg cells, only EWP feeding generated DORe BM-derived pTreg cells in mLNs, SI-LP, and SI-IE, and to lesser extents in popliteal LN and spleen, but not in the thymus (**Fig. 4B-4D**). Bulk mRNA-seq of DORe pTreg cells and WT nTreg cells in BMC mice having been EWP-fed for 8 weeks revealed the genes that were differentially expressed between the two Treg populations (**Fig. 4E**). Among such genes normalized for TPM (transcript per million) in the expression level, DORe-pTreg cells showed much higher expression of *Cd101* and *Gpr83* and lower expression of *Icos* when compared with WT nTreg cells from NC- or EWP-fed BMC mice (**Fig. 4F**). Regarding previously reported tTreg markers (Thornton et al., 2010; Weiss et al., 2012), *Ikzf2*(Helios) was low, whereas *Nrp1*(Neuropilin-1) was high, in DORe pTreg cells (**Fig. 4G**). Flowcytometric analysis further confirmed that DORe pTreg cells strongly expressed the CD101 protein in BMC mice (**Fig. 4H**). Similarly, in NC-fed WT (eFox) and DORe mice (**Fig. S2**), ∼80% of DORe pTreg cells and ∼25% of WT nTreg cells expressed CD101, while Foxp3^−^ Tconv cells scarcely expressed the molecule (**Fig. 4I**).

**Fig. 4.**
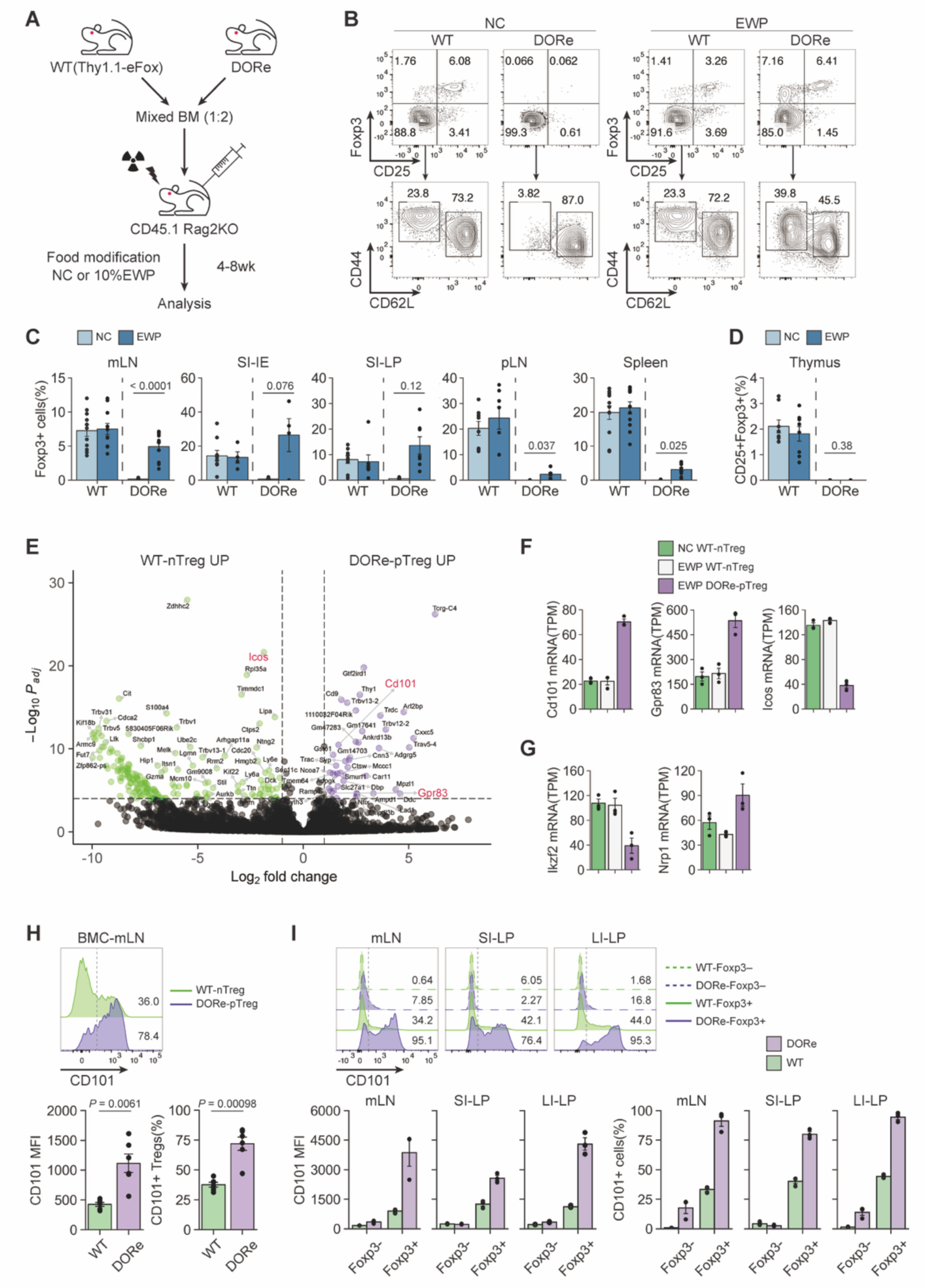
Specific expression of CD101 by pTreg cells generated by antigen feeding. (**A**) Irradiated CD45.1 Rag2KO mice reconstructed with CD3e^+^ cell-depleted bone marrow cells isolated from WT (Thy1.1-eFox) and DORe mice. Mixed BMC mice were analyzed at 4-8 weeks after BM cell transfer. (**B**-**D**) Representative flow cytometry profiles of WT- and DORe-derived CD4^+^ T cells in mLNs of BMC mice fed with NC- or EWP-chow for 4 weeks (B). Frequency of WT- and DORe-derived Foxp3^+^ cells in designated tissues (C) (*n* = 10-12) and in the thymus (*n* = 8) (D). (**E**) Volcano plot of differentially expressed genes between DORe-pTreg cells and WT-nTreg cells assessed by RNA-seq analysis. (**F, G**) TPM-normalized differentially expressed genes (F) and genes encoding known tTreg-markers (G) in BMC mice as shown in (B) (*n* = 3). (**H**) CD101 expression by DORe-pTreg cells and WT nTreg cells in BMC mice as shown in (B). Barplots indicate MFI of CD101 expression and frequency of CD101^+^ cells (*n* = 6). (**I**) CD101 expression by WT-Treg or Tconv cells, DORe-pTreg or Tconv cells in untreated mice. Barplots indicate MFI of CD101 expression and frequency of CD101^+^ cells (*n* = 3). Vertical bars indicate mean ± SEM. Statistical significance was assessed by unpaired *t* test or Welch’s *t* test (C, H).

To examine CD101 expression on Foxp3^+^ cells in a more physiological condition, DORe-derived naïve T cells were transferred into WT mice, which were then fed with EWP (**Fig. 5A**). Detectable numbers of DORe pTreg cells were found to be induced in mLN (**Fig. 5B** and **5C**). They highly expressed CD101 and Nrp-1 (**Fig. 5D**).

**Fig. 5.**
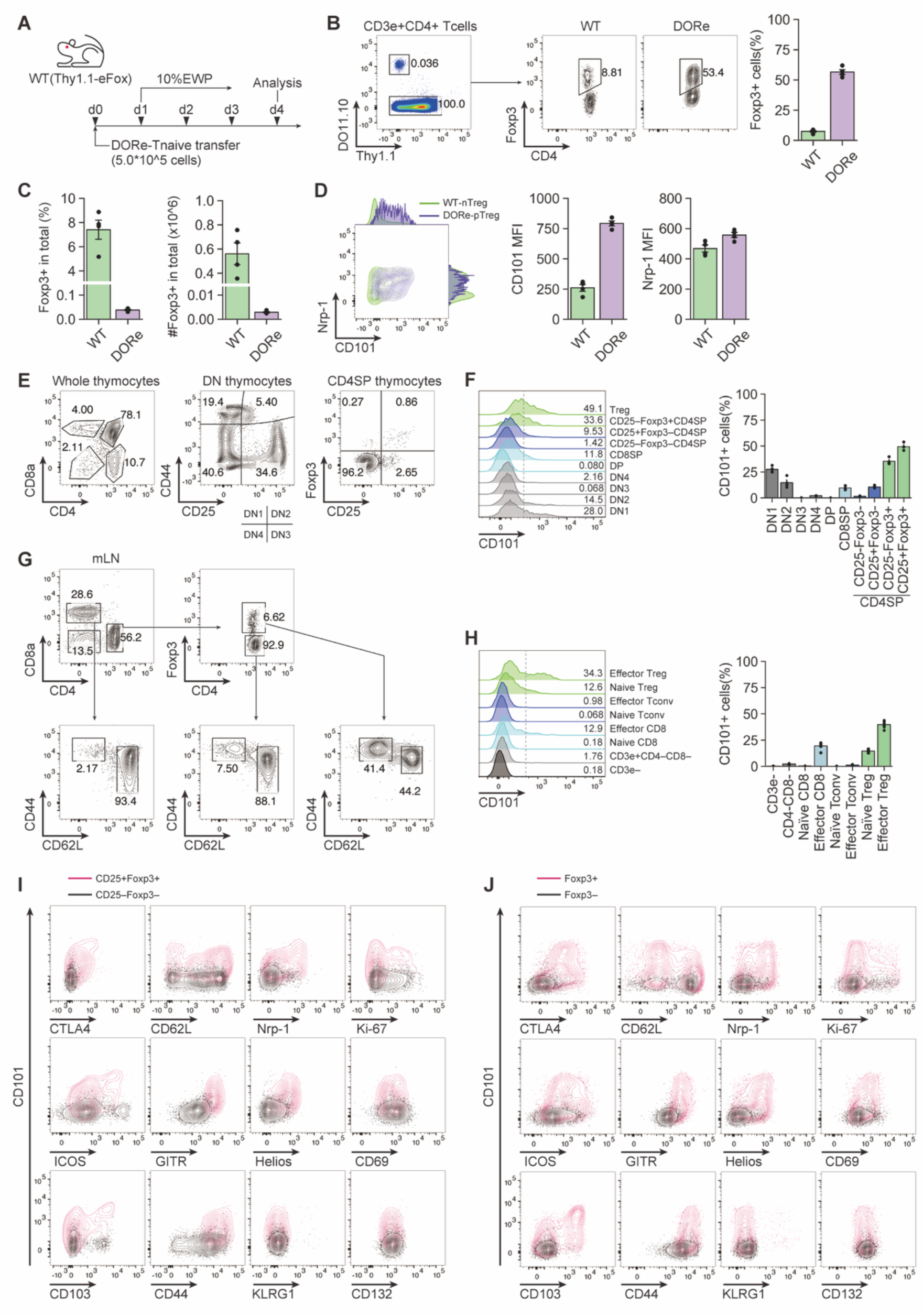
CD101 expression by pTreg cells, thymocytes, and peripheral T cells in WT mice. (**A**) Experimental scheme for assessing pTreg development in WT mice. Naïve CD4^+^ T cells from DORe mice were transferred into WT (Thy1.1-eFox) mice, which were then fed with EWP for 3 days. (**B**) Foxp3^+^CD4^+^ T cells in transferred DORe CD4^+^ T cells and in the host. Representative flow cytometry profiles of CD4^+^ T cells in mLNs of WT mice treated as shown in (A) and frequency of WT- and DORe-derived Foxp3^+^ cells (n = 4). (**C**) Frequencies and cell numbers of WT- and DORe-derived Treg cells in total CD4^+^ T cells. (**D**) CD101 and Nrp-1 expression by WT- and DORe-derived Foxp3^+^ cells. Representative flow cytometry profile and MFI of the expression (*n* = 4). (**E**) Gating strategy of thymocytes in WT mice. Whole thymocytes (left), CD4/CD8-double-negative (DN) thymocytes (middle), and CD4-single-positive (SP) thymocytes (right). (**F**) CD101 expression by thymocyte subpopulations. Representative histograms and frequencies of CD101^+^ cells in thymocyte fractions shown in (E) (*n* = 4). (**G**) Gating strategy of mLN-derived cells in WT mice. (**H**) Representative histogram of CD101 expression and frequencies of CD101^+^ cells in WT mLN-derived T-cell fractions (*n* = 6). (**I**, **J**) Representative flowcytometry profiles of CD101 and co-stained T cell-related molecules on Foxp3^+^ Treg or Foxp3^−^ T cells from the thymus (I) and mLNs (J). Vertical bars indicate mean ± SEM.

In the thymus of normal mice, CD101 was expressed by a fraction of DN1 (CD4/CD8 double-negative CD44^+^CD25^-^) thymocytes, losing its expression along differentiation and again becoming expressed at the CD25^−^Foxp3^+^CD4SP Treg precursor stage, with upregulation at mature CD25^+^Foxp3^+^ Treg cells (**Fig. 5E, 5F**). In the periphery, CD101 expression was limited to Foxp3^+^ T cells, with higher expression by CD44^+^CD62L^-^ effector Treg cells compared with naïve ones (**Fig. 5G, 5H**). In addition, CD101 expression by nTreg cells was well correlated with the expression of activation-associated molecules such as Nrp-1, GITR, CD69, CD103, and CD44, especially in mLNs (**Fig. 5I** and **5J**).

Taken together, these results indicate that intestinal pTreg cells generated by antigen feeding are distinct from tTreg cells in the expression levels of certain genes, especially CD101.

### *In vitro* induction of CD101^+^ iTreg cells from naïve and antigen-primed Tconv cells by blockade of costimulatory signal

Based on the above results, we investigated how functionally stable iTreg cells similar to CD101^+^ intestinal pTreg cells induced by antigen feeding could be generated *in vitro* from Tconv cells. Previous studies have shown that iTreg cells can be produced *in vitro* by anti-TCR stimulation with plate-bound anti-CD3 mAb in the presence of TGF-β, IL-2, and agonistic anti-CD28 mAb (Zheng et al., 2002; Chen et al., 2003), and that, in the presence of IL-2, iTreg generation was more efficient in the absence of agonistic anti-CD28 mAb (Mikami et al., 2020; Benson et al., 2007). Consistent with this, we found that iTreg cells generated without CD28 signal indeed expressed CD101 at higher levels than those produced with CD28 signal (**Fig. 6A** and **6B**). Moreover, CD28-deficient naïve Tconv cells more efficiently differentiated into iTreg cells with higher expression of CD101 compared with CD28-intact naïve Tconv cells; agonistic anti-CD28 mAb significantly hampered the generation of CD101^+^ iTreg cells from the latter, but not from the former (**Fig. 6C** and **6D**). Furthermore, TGF-β dose-dependent iTreg induction revealed that TGF-β significantly enhanced CD101 expression in a dose dependent manner only in iTreg cells generated without CD28 signal (**Fig. S4A**).

**Fig. 6.**
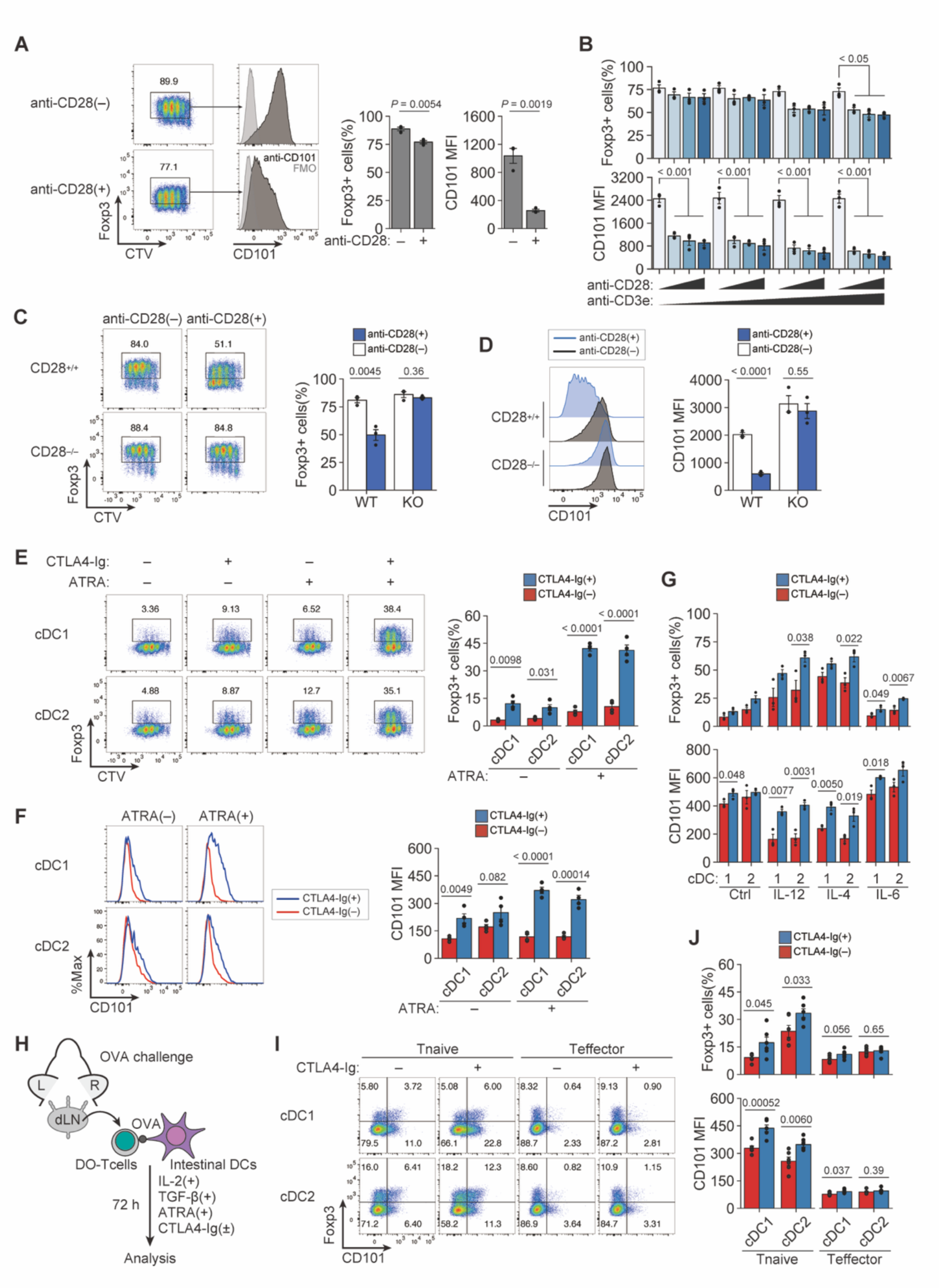
*In vitro* generation of CD101^+^ iTreg cells by absence or blockade of costimulatory signal during their induction. (**A**) *In vitro* generation of iTreg cells and their CD101 expression. iTreg cells were induced by plate-bound anti-CD3e mAb with or without anti-CD28 mAb in the presence of TGF-β and IL-2 (*n* = 3). (**B**) iTreg induction and their CD101 expression by anti-CD3e (0.5, 1, 5, and 10 µg/ml) and anti-CD28 mAb (0, 0.5, 1, and 2 µg/ml) stimulation as in (A) (*n* = 3). (**C**, **D**) iTreg induction from CD28-deficient or -intact CD4^+^ T cells (C) and their expression of CD101 (D) as in (A) (*n* = 3). (**E**, **F**) iTreg generation (E) and CD101 expression (F) by iTreg cells induced by co-culturing DO naïve Tconv cells with SI-LP-derived cDC1 or cDC2 in the presence of OVA and CTLA4-Ig (*n* = 4). (**G**) Effects of inflammatory cytokines (10 ng/ml each) and CTLA4-Ig on antigen-specific iTreg induction and their CD101 expression as in (E) and (F) (*n* = 3). 1 and 2 mean cDC1 and cDC2, respectively. (**H**) iTreg induction from antigen-specific Tconv cells in antigen-sensitized DORe mice. Naïve or effector Tconv cells from dLNs of antigen-sensitized DORe mice were co-cultured with SI-LP-derived cDC1 or cDC2 cells in the presence of OVA and CTLA4-Ig. (**I, J**) Frequencies of iTreg cells (I) and CD101 expression (J) of iTreg cells induced as shown in (H) (*n* = 6). Vertical bars indicate mean ± SEM. Statistical significance was assessed by Tukey’s HSD test (B) and unpaired *t* test or Welch’s *t* test (C, D, E, F, G, J).

Next, to simulate *in vitro* the condition of intestinal pTreg generation in EWP-fed DORe mice, we examined the effects of CD80/CD86 blockade with CTLA-4-immunoglobulin (CTLA4-Ig) fusion protein on iTreg generation from naïve DORe Tconv cells stimulated with OVA in the presence of splenic DCs (**Fig. S4B**). CTLA4-Ig did not inhibit proliferative differentiation of Foxp3^+^ cells even at high doses (**Fig. S4C**), increased CD101 expression levels in an IL-2 dependent fashion (**Fig. S4D**), and conferred *Foxp3*-CNS2 hypomethylation on the generated iTreg cells (**Fig. S4E**). With cDC1 and cDC2 cells prepared from SI-LP (**Fig. S4F**), CTLA4-Ig in combination with all-trans retinoic acid (ATRA) significantly increased the generation of Foxp3^+^ cells and enhanced their expression of CD101 on both types of DCs (**Fig. 6E** and **6F**). CTLA4-Ig together with ATRA increased iTreg generation and CD101 expression even in the presence of various inflammatory cytokines (**Fig. 6G**). Furthermore, with naïve or effector/memory Tconv cells from dLNs of OVA immunized DORe mice (**Fig. 6H**), CTLA4-Ig effectively augmented iTreg generation and their CD101 expression with naïve Tconv cells and, to lesser degrees, with effector/memory Tconv cells (**Fig. 6I** and **6J**).

Taken together, blockade of CD80/CD86 did not hamper iTreg generation with TGF-β, IL-2, and antigen stimulation, rather enhancing the generation of iTreg cells, CD101^+^ ones in particular. Thus, CD28 signaling is dispensable for iTreg generation, especially in the presence of IL-2. CD101 expression could be attributed to the reduction of CD28 co-stimulatory signal in iTreg generation.

### Establishment of oral tolerance in antigen-presensitized mice by CD80/CD86 blockade and subsequent antigen feeding

The above result that CTLA4-Ig enabled iTreg cell induction from antigen-primed Tconv cells prompted us to determine whether *in vivo* CTLA4-Ig treatment would be able to establish oral tolerance even in precedingly antigen-primed mice (**Fig. 7A**). When DORe mice were OVA sensitized at the right ear, injected intraperitoneally (i.p.) with CTLA4-Ig at various doses or saline, and then fed with EWP chow for 4 days, the right ear swelling gradually subsided during the feeding whether the mice were treated with CTLA4-Ig or not (**Fig. S5A**). However, mice treated with saline or low-dose (2 µg) CTLA4-Ig showed significant weight loss during EWP feeding (**Fig. S5B**), contrasting with no weight loss in the mice CTLA4-Ig-treated at high (10 or 50 µg) doses. The weight loss recovered after cessation of the feeding (**Fig. 7B**). When these mice were OVA challenged to the left ear after 4-day EWP feeding, those treated with high dose (10 or 50 µg) CTLA4-Ig did not show ear swelling or succumb to histologically evident inflammation, contrasting with other groups suffering from severe ear swelling and dermatitis (**Fig. 7C** and **S5C**). Notably, contrasting with the CTLA4-Ig/EWP-treated mice exhibiting no ear swelling, the CTLA4-Ig/NC-treated mice developed severe ear swelling and dermatitis. The prevention of ear swelling was also observed with the higher antigen dose (30% EWP-containing chow), while the lower antigen dose (1% EWP) displayed a reduced preventive effect (**Fig. S5D-F**). These results taken together indicated that both CTLA4-Ig treatment and antigen feeding were required for inducing the unresponsiveness.

**Fig. 7.**
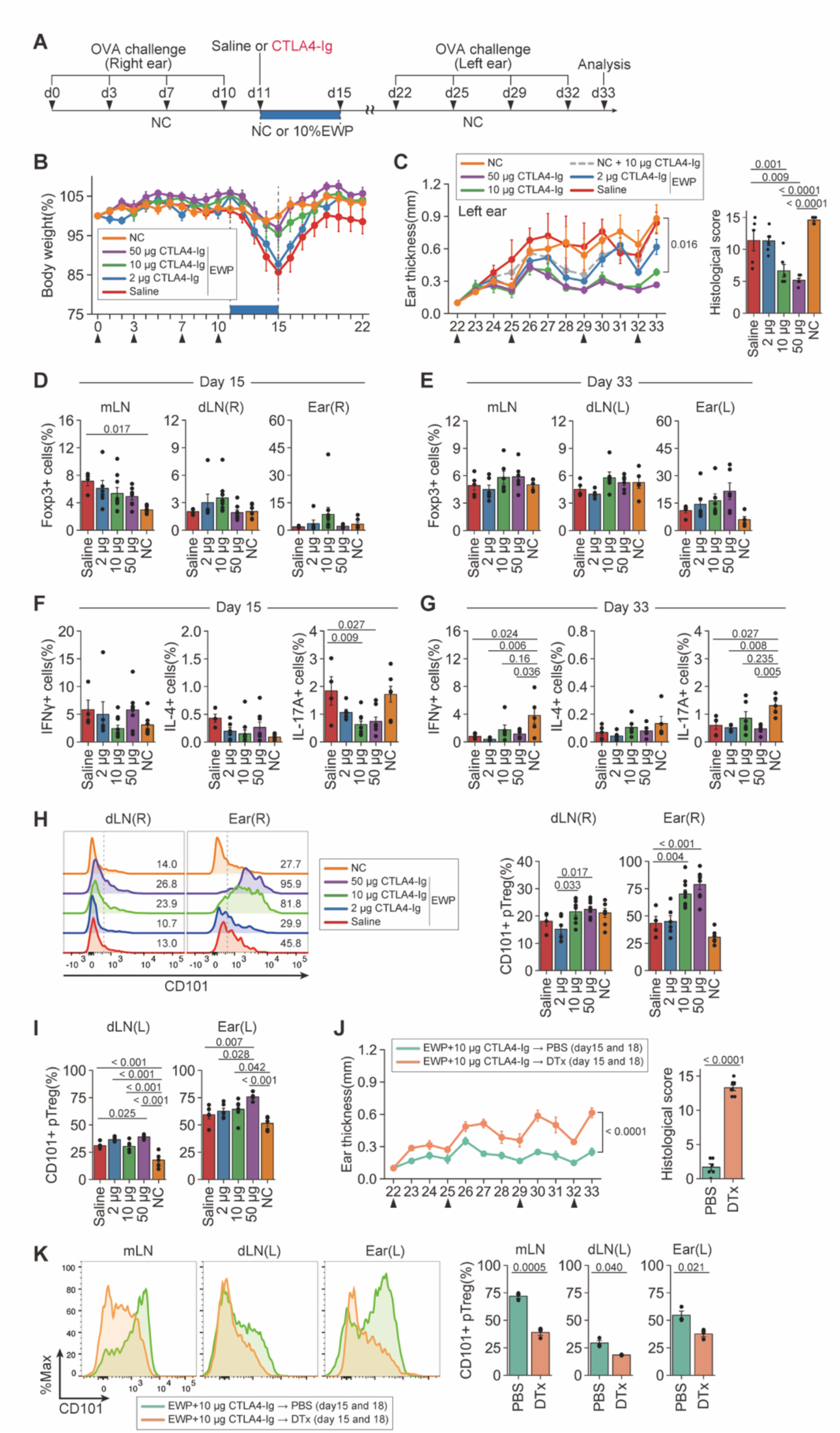
Establishment of oral tolerance in antigen-presensitized mice by CD80/CD86 blockade and subsequent antigen feeding. (**A**) Experimental protocol. DORe mice were OVA sensitized at the right ear four times, i.p. injected with CTLA4-Ig or saline on day 11, fed with EWP chow or NC for four days, OVA challenged at the left ear four times from day 22, and assessed for ear swelling. (**B, C**) Changes in body weight (B), left ear thickness and histological scores (C) (*n* = 5-6). Arrowheads indicate OVA challenges. (**D**, **E**) Frequencies of pTreg cells in mLNs, dLNs, and the right ear at day 15 (D) and day 33 (E) (*n* = 5-6). (**F**, **G**) Frequencies of IFN-γ^+^, IL-4^+^, and IL-17A^+^ cells in dLNs at day 15 (F) and day 33 (G) (*n* = 5-6). (**H**) CD101 expression by pTreg cells in dLNs and the right ear at day 15. Numbers in the histogram are representative of the frequency of CD101^+^ cells. Barplots indicate the frequency of CD101^+^ pTreg cells (*n* = 4-10/group). (**I**) Frequencies of CD101^+^ cells within pTreg cells in dLNs and the left ear at day 33 (*n* = 5-6). (**J**, **K**) Effects of Treg depletion after CTLA4-Ig injection and EWP feeding, After CTLA4-Ig injection and EWP feeding, DORF mice were i.p. treated with PBS or DTx at days 15 and 18, and re-challenged with OVA (arrowheads). Changes in left ear thickness at day 33 in DORF mice (J) (*n* = 6-7). CD101 expression by pTreg cells in mLNs, dLNs, and the left ear (K). Barplots indicate the frequencies of CD101^+^ pTreg cells (*n* = 3). Vertical bars indicate mean ± SEM. Statistical significance was assessed by Tukey-Kramer test (C, D, E, F, G, H, I) and unpaired *t* test or Welch’s *t* test (J, K).

There were little differences among these mouse groups in the ratio of Foxp3 cells on day 15 and 33, with a tendency of Treg increase in the CTLA4-Ig-treated mice (**Fig. 7D** and **7E**). The ratios of cytokine-secreting cells among DO Tconv cells were not significantly different among the mouse groups at day 15, except significant reduction of the ratio of IL-17^+^ cells in dLNs of the CTLA4-Ig-treated groups compared with the saline-treated or NC-fed mice (**Fig. 7F**). IFN-γ^+^ cells as well as IL-17^+^ cells in the former were significantly lower compared with the latter at day 33 (**Fig. 7G**). The results suggested that the high ratios of Foxp3^+^ cells in low-dose CTLA4-Ig- or saline-treated group could be attributed to their secondary expansion due to inflammation.

The pTreg cells having developed in EWP-fed mice, whether treated with CTLA4-Ig or saline, showed Treg-specific DNA demethylation at the *Foxp3* CNS2 region, suggesting their functional stability as Treg cells (**Fig. S5G**). In contrast to no significant differences in total Foxp3^+^ cells in the ear or dLNs (**Fig. 7D** and **7E**), CD101 expression by Foxp3^+^ cells was significantly upregulated and the ratio of CD101^+^Foxp3^+^ cells increased at day 15 in the right ear and its dLNs of the CTLA4-Ig-treated mice depending on the dose of CTLA4-Ig, similarly in the left ear and its dLNs at day 33 (**Fig. 7H** and **7I**). Depletion of Foxp3^+^ cells by DTx administration on day 15 and 18 in CTLA4-Ig-treated and then EWP-fed DORF mice resulted in an exacerbated ear inflammation, a decrease in pTreg cells in the inflamed ear tissue, and an increase in IL-4^+^ T cells in dLNs (**Fig. 7J, S5H**, and **S5I**). CD101^+^ Foxp3^+^ cells were also significantly decreased in the inflamed ear tissue, dLNs and mLNs (**Fig. 7K**).

These results collectively demonstrate that CD80/CD86 blockade by CTLA4-Ig and subsequent antigen feeding is able to establish oral tolerance in antigen-presensitized mice through inducing CD101^+^ pTreg cells.

## Discussion

It has been controversial whether pTreg cells are as functionally stable as tTreg cells and how they can be functionally and phenotypically distinguished from the latter. We have shown here that intestinal pTreg cells induced by antigen feeding are similar to tTreg cells. They not only express Foxp3 but also possess Treg-type epigenetic changes such as DNA hypomethylation at the Treg-specific demethylation regions and activation of the Treg-specific super-enhancers at Treg signature gene loci including *Foxp3*, *Il2ra*, and *Ctla4*. Additionally, these cells are Helios^hi^ and Neuropilin-1^hi^ as are tTreg cells (Thornton et al., 2010; Weiss et al., 2012), while pTreg cells newly generated in BMC mice by EWP feeding were Helios^low^. It is likely that chronic stimulation may gradually upregulate Helios and Neuropirin-1 in pTreg cells (Szurek et al., 2015; Thornton and Shevach, 2019). In addition, the generated pTreg cells are in a tissue-adapted state as seen with tissue-specific and -resident tTreg cells (Muñoz-Rojas and Mathis, 2021; Dikiy and Rudensky, 2023), as illustrated by the expression of *Tnfrsf9*, *Tnfrsf4*, *Gata3*, and *Areg* as well as skin-specific chemokine receptors such as *Ccr5* and *Ccr8* (Schiering et al., 2014; Miragaia et al., 2019). However, despite these immunological properties indicative of functional competence and stability of pTreg cells induced by antigen feeding, cessation of antigen administration resulted in decline of oral tolerance with diminution of proliferative and Bcl-2^+^ (i.e., apoptosis resistant) pTreg cells. This could be, at least in part, attributed to the antigen specificity of the induced pTreg cells, as well as the strength and duration of antigen stimulation delivered to them in the periphery. tTreg cells are more self-reactive than Tconv cells, hence tonically stimulated in the periphery for better survival (Hsieh et al., 2004, 2012). In contrast, pTreg cells once generated by antigen feeding may require persistent or frequent stimulation by the administered cognate non-self antigen for their survival to maintain tolerance, despite their possession of Treg-specific epigenetic alterations and acquisition of tTreg-like immunological properties.

The generation of intestinal pTreg cells by antigen feeding depends on the quantity of cognate antigen contained in the diet. Notably, even DORe mice fed with NC or Ag-free diet developed Foxp3^+^ T cells particularly in SI-LP and LI-LP, and that they are phenotypically and epigenetically indistinguishable from pTreg cells generated by EWP-high diet. In NC-fed mice, a certain diet protein(s) or a microbial substance(s) cross-reactive with OVA could likely stimulate DO Tconv cells to differentiate into pTreg cells. The NC-induced pTreg cells were, however, smaller in number and less activated (e.g., lower in CD25 expression) than the high-EWP-induced pTreg cells and unable to suppress ear swelling upon antigen exposure. These findings indicate that induction of oral tolerance depends on the number of antigen-specific pTreg cells and/or their activation status, and that both events depend on the quantity of the ingested antigen. This dependency of oral tolerance on the quantity of administered antigen may also be the case in establishing mucosal tolerance via other routes, for example, by sublingual antigen administration for allergy treatment (Windom et al., 2024).

Intestinal pTreg cells induced by antigen feeding specifically express CD101, which is also highly specific for iTreg cells generated *in vitro* by the absence or blockade of CD28 co-stimulatory signal. Intestinal pTreg cells found in NC-fed mice, as discussed above, similarly expressed CD101, whereas nTreg cells in mLNs in WT mice hardly expressed the molecule even after high-EWP feeding. Furthermore, *in vivo* treatment with CTLA4-Ig for oral tolerance induction generated CD101^+^ pTreg cells in the antigen-challenged tissue, and the degree of expression depended on the dose of CTLA4-Ig. Recent studies have shown that CD101 is highly expressed by tissue-resident Treg cells compared to circulating ones (Kaminski et al., 2023); its expression is correlated with the level of FOXP3 expression in human Treg cells (Ferraro et al., 2014), and with *in vivo* Treg suppressive capacity on GvHD and IBD in mouse models (Fernandez et al., 2007; Schey et al., 2016). Furthermore, a genetic variant of CD101 contributes to genetic susceptibility to type 1 diabetes in an animal model through affecting Treg and other immune cells (Rainbow et al., 2011; Mattner et al., 2019). CD101 is reportedly expressed by exhausted CD8^+^ T cells as well (Hudson et al., 2019). These findings and ours, when taken together, suggest that CD101 plays a role in modulating TCR and CD28 signaling (Soares et al., 1998), especially the latter, for pTreg development in the intestine and iTreg generation from Tconv cells *in vitro*. The molecular basis of the function of CD101 and its ligand, which is currently unknown, remains to be determined. Nevertheless, CD101 could be a useful marker for pTreg cells, especially those induced by CD80/CD86^low^ tolerogenic DCs, which transduce TCR signal with reduced CD28 signal.

CD80/CD86 blockade by CTLA4-Ig facilitates *in vivo* CD101^+^ pTreg induction and *in vitro* CD101^+^ iTreg generation. The blockade and antigen stimulation allow both Treg populations to proliferate and enable them to acquire the Treg-type epigenome, hence their functional and cell-lineage stability (Mikami et al., 2020). IL-2 is essential for this Treg generation because it can compensate for the absence or reduction of CD28 signaling, hampering anergy induction and enabling more efficient generation of iTreg cells than without the blockade. It is plausible that *in vivo* provision of IL-2 could further facilitate pTreg cell generation by antigen feeding after CD80/CD86 blockade. In addition, APCs, especially DCs, play key roles in pTreg cell induction by antigen feeding (Esterházy et al., 2016; Brown and Rudensky, 2023). Both cDC1 and cDC2 were able to generate iTreg cells *in vitro* under CD80/CD86 blockade. In addition to the role of CD80/CD86 for Treg induction by APCs, it has been shown that PD-L1^+^ DCs are potent in inducing antigen-specific Treg cells and that PD-L1 blockade inhibits the induction (Wang et al., 2008; Francisco et al., 2009). We and others have shown that Treg cells are able to inhibit the T-cell stimulatory activity of APCs by reducing their CD80/CD86 expression via trogocytosis and transendocytosis, which are dependent on Treg-expressed CTLA-4 (Wing et al., 2008; Onishi et al., 2008; Qureshi et al., 2011; Tekguc et al., 2021). This CD80/CD86 reduction on APCs results in dissociation of PD-L1 from *cis*-bound CD80, increasing free-PD-L1 available for the inhibition of PD-1 expressing effector T cells (Tekguc et al., 2021). CTLA4-Ig can have a similar effect by disrupting *cis*-CD80-PD-L1 interaction depending on the ratio of CD80 to PD-L1 on APCs (Tekguc et al., 2021; Oxley et al., 2024; Robinson et al., 2024). We have shown in this report that reduction or abrogation of CD28 signal alone (for example, by utilizing CD28-deficient naïve Tconv cells), without modulating PD-L1/PD-1 interaction, suffices to generate iTreg cells effectively. It still needs to be determined, however, whether enhanced signaling from free-PD-L1 on APCs to PD-1^+^ Tconv cells facilitates the conversion of the latter to pTreg cells.

Prior antigen sensitization and the following antigen feeding evoked systemic inflammation. CTLA4-Ig treatment before antigen feeding not only prevented the systemic inflammation but also suppressed local inflammation at the site of antigen exposure. These findings carry some clinical implications. First, it is conceivable that certain food allergies might be a consequence of prior antigen sensitization at other sites of the body, as proposed as the dual-allergen-exposure hypothesis for food allergy (Lack, 2008). Second, CTLA4-Ig has been used for treatment of allergy on the assumption that it may suppress effector T-cell activation and IgE production (Van Wijk et al., 2005). We have shown here that CTLA4-Ig can additionally facilitate the generation of antigen-specific functionally stable pTreg cells from antigen-primed Tconv cells when combined with subsequent antigen feeding. Third, this approach to establishing oral tolerance by CD80/CD86 blockade and antigen feeding can be useful for mucosal tolerance induced by other ways, for example, by sublingual or subcutaneous antigen treatment to treat chronic allergy. It could also be extended to the induction of immune tolerance towards microbial antigens derived from intestinal commensal bacteria to treat inflammatory bowel disease.

In conclusion, antigen-specific functionally stable CD101^+^ pTreg cells are generated in the intestine by antigen feeding and engaged in the establishment of long-term systemic antigen-specific immune tolerance. CD101^+^ iTreg cells with similar phenotype and function can also be produced *in vitro* from antigen-specific Tconv cells by depriving CD28 costimulation during their induction, suggesting a common developmental basis for these pTreg and iTreg cells. Furthermore, continuous antigen feeding combined with CD28 co-stimulation blockade is able to generate CD101^+^ pTreg cells and establish immune tolerance even in previously antigen-sensitized animals.

## Materials and methods

### Mice

BALB/c mice were purchased from SLC or CLEA. DO11.10 TCR transgenic mice, *Rag2*-deficient mice, Foxp3-eGFP (eFox) reporter mice, Thy1.1-eFox mice, Foxp3-DTR-GFP mice, and *CD28*-deficient mice were previously described (Mikami et al., 2020; Kim et al., 2007; Shahinian et al., 1993). All mice were maintained under specific pathogen-free conditions. All procedures were conducted in accordance with the National Institutes of Health Guide for the Care and Use of Laboratory Animals and approved by the Committee on Animal Research of Osaka University.

### Food modification

AIN-93G diet and egg-white-powder (EWP)-added food were purchased from CLEA. AIN-93G diet was used as antigen-free food (**Table. 1**). CE-2 (CLEA) as normal chow (NC) was used as standard food in the animal facility. For DORe mice, food modification began at 2 weeks of age under milk feeding or at 4 weeks of age after weaning, and continued until 8 weeks of age. For bone marrow chimeric mice, the food modification was started at the time of bone marrow cell transfer, and continued for 4-8 weeks.

### Epicutaneous sensitization of OVA antigen to mice

The procedure was described in previous studies (Shimura et al., 2016). Mice were anesthetized with isoflurane. Both ears were treated with tape stripping five or six times using surgical tape (21N, Nichiban, Tokyo, Japan). Then, a total of 12.5 µl of 10 mg/ml of papain with 10 mg/ml OVA was applied to each side of the surface of mouse ear. This treatment was repeated twice per week for 2 weeks. During the experimental period, ear thickness was measured daily using a digital caliper. For depleting Foxp3^+^ cells in the experiment, diphtheria toxin (400 ng in 200 µl PBS) was injected intraperitoneally twice per week into the Foxp3-DTR-GFP background mice.

### Histopathology

Tissues were fixed with 10% formalin, embedded in paraffin blocks, and sectioned and stained by hematoxylin and eosin. For ear swelling test, histological and pathological scoring was performed by following criteria and summed up; disorder of epidermis: 1-3, disorder of dermis: 1-3, disorder of hypodermis: 1-3, cell invasion: 1-3, and range of inflammation: 1-3.

### Isolation of ear-infiltrating immune cells

Dissected ears were minced into small pieces with scissors and digested in the isolation buffer (HBSS(−) with 2% FBS, 2 mg/ml Collagenase D (Roche), and 20 µg/ml DNaseI (Roche)) with 1,300 rpm shaking at 37℃ for 2 h. The suspension was then passed through a 70-μm mesh and mash the debris by using a syringe plunger. The flow-through was centrifuged at 500xg at 4℃ for 5 min. The pellet was washed and resuspended with FACS buffer. Isolated cells were used for flowcytometry directly or first stimulated with PMA and ionomycin in the presence of GolgiStop (BD) for 4 h at 37℃ for subsequent intracellular cytokine staining.

### Isolation of lymphocytes from intestinal mucosa

Intestines were harvested from mice, the lumen was opened longitudinally with scissors, the luminal contents were removed, and the tissues were washed with FACS buffer. After cutting the tissues into 2 cm, they were incubated with IE isolation buffer (HBSS(−) with 2% fetal bovine serum (FBS), 1 mM EDTA (Nacalai)) at 37℃ for 30 min. The tissues were then incubated with LP isolation buffer (HBSS(−) with 2% FBS, 1 mg/ml collagenase D (Roche), and 10 µg/ml DNaseI (Roche)) at 37℃ for 30 minutes. The suspension was then passed through a 70-μm mesh, and the debris was mashed using a syringe plunger. The flow-through was centrifuged at 500xg at 4℃ for 5 min. The pellet was resuspended with 40% Percoll solution, then layered with 80% Percoll solution and centrifuged at 2,000 rpm at room temperature for 20 min without acceleration and brake. The interface was collected and washed with FACS buffer.

### Generation of mixed bone marrow chimeric mice

Bone marrow cells were harvested from the femur and tibia of mice, using a 23G needle and syringe. After lysing red blood cells, CD3e^+^ T cells were removed using the MACS negative selection system (Miltenyi Biotec) or the BD IMag Cell Separation System (BD) following the manufacturer’s protocol. T cell-depleted bone marrow cells from Thy1.1-eFox and DORe mice were mixed at a ratio of 1:2 in PBS buffer and injected intravenously into CD45.1^+^ *Rag2*-deficient mice as recipientsthat had been irradiated at 4.0 Gy. After 4 weeks or more later from bone marrow reconstitution, chimeric mice were sacrificed and analyzed.

### Cell preparation, cell sorting, and flowcytometry

For the preparation of DORe-pTreg cells, CD4^+^DO11.10^+^CD25^+^Foxp3-eGFP^+^ cells were sorted from the peripheral lymph nodes and peripheral tissues harvested from DORe mice. Peripheral Tconv cells were defined as these gating below; naïve Tconv cells (CD4^+^CD25^−^Foxp3-eGFP^−^CD62L^hi^CD44^lo^) and effector Tconv cells (CD4^+^CD25^−^Foxp3-eGFP^−^CD62L^lo^CD44^hi^). As antigen presenting cells, cDC1 (CD45^+^CD11c^+^IA-IE^+^CD103^+^CD11b^−^) and cDC2 (CD45^+^CD11c^+^IA-IE^+^CD103^+^CD11b^+^) fractions were sorted from SI-LP in CD45.1 WT mice. BD Cytofix/Cytoperm Fixation/Permeabilization Kit (BD) was used for intracellular staining for cytokines used following stimulation with PMA and Ionomycin and treatment with Golgi-Stop (BD Biosciences), and the Foxp3 staining kit (eBiosciences) was used for transcriptional factors or intracellular molecules. Flowcytometry analysis and cell sorting were performed using FACSCantoII, FACSCellesta, FACSAriaII, and FACSAriaFusion (BD Biosciences). Antibodies used in this study are listed (**Table S2**).

### CpG methylation analysis by bisulfite sequencing in Treg-specific demethylation regions

Bisulfite sequencing analysis was performed as previously described (Arai et al., 2022). Briefly, cells were sorted by FACSAriaII, and gDNA was extracted by phenol extraction followed by ethanol precipitation. Bisulfite base conversion was carried out using the Methyl Easy Xceed Rapid DNA Bisulphite Modification Kit (Human Genetic Signatures) or following our original methods previously described. PCR primer sequences for Treg-specific demethylated regions are available (Ohkura et al., 2012; Arai et al., 2022).

### ChIP-seq

ChIP-seq experiments were performed as previously described (Kitagawa et al., 2017; Kawakami et al., 2021). Briefly, sorted cells were fixed with 1% formaldehyde (ThermoScientific) for 10 min for anti-histone ChIP at room temperature. After nuclear extraction, chromatin lysate was fragmentated using Picoruptor (Diagenode), 30 sec sonication and 30 sec cooling for 7 cycles at 4℃, before immunoprecipitation. Immunoprecipitated chromatin lysate was reverse cross-linked at 65℃ for 24 h, followed by purification and library preparation using NGS Library Preparation kit for IonS5 (ThermoScientific) or NEBNext Ultra II DNA Library Prep Kit for Illumina (Illumina) according to the manufacturer’s instructions. Raw data were generated using the IonS5 sequencing system (ThermoScientific) or NextSeq500 (Illumina).

### ATAC-seq

ATAC-seq was performed as previously described (Corces et al., 2017). Briefly, sorted cells (up to 100,000 cells) were lysed with100 µl of lysis buffer (0.01% digitonin, 0.1% NP-40, 0.1% Tween 20 in resuspension buffer; 10 mM Tris-HCl pH7.5, 100 mM NaCl, 3 mM MgCl_2_) for 3 min on ice. After removal of lysis buffer by centrifugation, Tn5 tagmentation was performed using Illumina Tagment DNA TDE1 Enzyme and Buffer Kits (Illumina) at 37℃ for 30 min, with shaking at 1,000 rpm. After purification using DNA Clean & Concentrator-5 (Zymo Research), tagmented DNA was amplified using NEBNext High-Fidelity PCR Master Mix (New England BioLabs) with the following prepared DNA libraries were purified using DNA Clean & Concentrator-5 or Ampure XP (Beckman Coulter). Sequencing was performed using NextSeq500 or NovaSeq (Illumina).

### Data processing and analyses of ChIP-seq and ATAC-seq data

For ChIP-seq analysis, raw sequence reads were inspected and trimmed using Trim-galore (v0.6.6) (Babraham Bioinformatics). Trimmed reads were mapped to mm10 genome using Bowtie2 (v2.3.5) (Langmead and Salzberg, 2012) with the following options; --local --very-sensitive-local. Mapped reads were sorted using Samtools (v1.6)(Li et al., 2009), then converted into bigwig files for visualization by bamCoverage (v3.1.3), included deepTools package(Ramírez et al., 2016), with the following options; - of bigwig --binSize 5 -p max --normalizeUsing CPM --smoothLength 15 –ignoreDuplicates. Peak calling was performed using MACS2 (v2.1.1.20160309)(Feng et al., 2012) with the following options; macs2 callpeak -t ${filename} -name ${outputfilename} -f BAM -g mm --SPMR --nolambda –nomodel --bdg --call-summits. For defining Treg or Tconv-specific enhancer regions, narrowPeak files were concatenated, sorted. Sorted narrowPeak file was merged using bedtools (v2.30.0)(Quinlan and Hall, 2010) and then converted into SAF format file. SAF peak file was annotated using HOMER (v4.11)(Heinz et al., 2010). Count files were generated using featureCounts(Liao et al., 2014) with the following options; -a annotated.saf -F SAF -p -o counts.txt ${BAM}. Differential expression analysis was performed using DESeq2. To assess the tag density of H3K27ac signals in Treg and naïve Tconv cells, matrix files were created using computeMatrix (v3.1.3), included deepTools package, with the following options; --beforeRegionStartLength 2000 --regionBodyLength 5000 --afterRegionStartLength 2000 --skipZeros, then visualized plotProfile command.

For ATAC-seq analysis, sequence reads were mapped to mm10 genome using Bowtie2 (v2.3.5) by following options; --very-sensitive. Mapped reads were sorted by Samtools (v1.9) and then converted into bigwig files for visualization by bamCoverage (v3.1.3) with the following options; -of bigwig -- binSize 5 -p max --normalizeUsing CPM --smoothLength 15 –ignoreDuplicates. Peak calling was performed using MACS2 (v2.1.1.20160309) with the following options; macs2 callpeak -t ${filename} - name ${outputfilename} -f BAM -g mm --SPMR --nolambda --nomodel --shift -75 --extsize 150 --keep- dup all --bdg --call-summits.

IGV Integrated genome viewer was used to visualize peak or region data.

### Single-cell RNA-seq and analysis

Single-cell RNA-Seq was performed on a Chromium instrument (10X Genomics) following the user guide manual for 30 v3 or 50 v2 chemistry. Briefly, FACS-sorted DORe-derived T cells isolated from mLNs and SI-LP were stained with TotalSeq anti-mouse Hashtag Antibody (BioLegend), and washed three times with FACS buffer. Cells were resuspended with FACS buffer to a final concentration of approximately 1,000 cells/µl with a viability above 90%. Capturing cells in droplets, reverse transcription, and cell barcoding were performed on the Chromium Controller (10x Genomics), followed by PCR amplification and library construction using Chromium Next GEM Single Cell 5’ Kit v2. Final libraries were sequenced on NovaSeq6000 (Illumina), 20,000 reads/cell for mRNA and 5,000 reads/cell for antibody-derived tags.

Sequenced reads were quantified using Cell Ranger (v5.0.0) with the pre-built reference refdata-gex-mm10-2020-A downloaded from 10x Genomics’ website. Quantified expressions were preprocessed and visualized using Seurat (v4.1.3 or v5.0.1)(Hao et al., 2021). Differentially expressed genes were calculated using “FindMarkers” function based on the non-parametric Wilcoxon rank sum test in Seurat. Gene ontology analysis was performed using clusterProfiler (v4.10.0)(Wu et al., 2021). For the trajectory analysis, selected cells were re-clustered and ordered using Monocle3 (v1.3.4)(Cao et al., 2019).

### Bulk RNA-seq and analysis

Bulk RNA-seq was performed as below. Briefly, FACS-sorted cells were lysed by RLT RNA lysis buffer (Qiagen) containing 2-mercaptoethanol to a final concentration of approximately 1,000 cells/10 µl followed by reverse transcription using the SMART-seq v4 Ultra Low Input RNA Kit for Sequencing (Clontech). Library preparation was performed using the Kapa Library preparation kit for IonTorrent or for Illumina (KAPA) by following the manufacturer’s protocol. Sequencing of cDNA libraries was performed on IonS5 (Thermo Scientific), NextSeq500, or NovaSeq6000 (Illumina).

Sequenced reads were inspected and trimmed using Trim-galore (v0.6.6) and aligned to the mouse genome GRCm38/mm10 using STAR (v2.7.7a)(Dobin et al., 2013). Reads counts at the gene level were calculated using RSEM (v1.3.1)(Li and Dewey, 2011). Normalization for library size and differential expression analysis were performed using DESeq2 (v1.32.0)(Love et al., 2014).

### Cell culture

Cell culture was performed in RPMI 1640 supplemented with 10%(v/v) FBS, 60 g/ml penicillin G, 100 g/ml streptomycin, and 0.1 mM 2-mercaptoethanol. For in vitro Treg induction, sorted naïve Tconv cells were stimulated with plate-bound anti-CD3e mAb (clone: 145-2C11) and soluble anti-CD28 mAb (clone: 37.51) in the presence of 100 U/ml hIL-2 (Shionogi) and 5 ng/ml rhTGF-β_1_ (R&D Systems). For the co-culture experiment, DORe-derived naïve or effector/memory Tconv cells were stimulated by sorted APCs, 5 µM OVA_323-339_ peptide (MBL Co., LTD., MedChemExpress), 100 U/ml rhIL-2, and 2.5 ng/ml rhTGF-β_1_ and with or without 10 nM ATRA supplementation (Wako) or 10 µg/ml CTLA4-Ig (abatacept; Orencia, BMS) for 72 hours.

### Statistics

Values were expressed as mean ± SEM. Statistical significance was assessed by Student’s *t* test or Welch’s *t* test (two groups) following *F* test. One-way ANOVA and Tukey’s HSD or Tukey-Kramer test were conducted for multiple comparisons. A *p*-value < 0.05 was considered statistically significant, and every *p* value was shown in the figures.

## Online supplemental materials

Fig. S1 shows pTreg generation and maintenance by antigen feeding. Fig. S2 shows the development of pTreg cells in NC-fed DORe mice. Fig. S3 shows single-cell RNA-seq analysis of T cells in DORe mice and bulk mRNA-seq analysis of pTreg cells from different tissues after EWP feeding. Fig. S4 shows dose effects of CTLA-4-Ig, TGF-β, and IL-2 on iTreg generation and the gating strategy for isolating intestinal DCs. Fig. S5 shows the effects of CD80/CD86 blockade and subsequent antigen feeding on antigen-presensitized mice. Table S1 indicates nutritional facts of modified foods. Table S2 lists the antibodies used in the study.

## Data availability

The raw data (fastq files) from ATAC-seq, ChIP-seq, RNA-seq, and scRNA-seq experiments have been deposited in the DDBJ BioProject database, with links to BioProject accession number PRJDB18773. All data needed to evaluate the conclusions are present in the paper or the Supplementary Materials.

## Acknowledgments

We thank all Sakaguchi lab members, particularly: R. Ishii and M. Matsuura for their technical assistance, C. Tay and S.K. Singh for critical reading of the manuscript. We would like to thank K. Kawade for advising us about the production of modified foods. The genomic sequencing and bioinformatic analysis were conducted using the facility at the Genome Information Research Center of the Research Institute for Microbial Diseases, Osaka University.

## Funding

This study was supported by the Japan Society for the Promotion of Science (JSPS) grant-in-aid for JSPS Fellows (20J20443 to M.A.), JSPS Grant-in-Aid for Early-Career Scientists (20K16279 to R.K.), JSPS Grant-in-Aid for Scientific Research (23H02733 to R.K.), INFRONT Office of Director’s Research Grants Program (to R.K.), Grant from Nipponham Foundation for the Future of Food (to R.K.), JSPS Grant-in-Aid for Specially Promoted Research (16H06295 to S.S.), JSPS Grant-in-Aid for Scientific Research (24H00610 to S.S.), and Leading Advanced Projects for Medical Innovation (LEAP; 18gm0010005 to S.S.).

## Author contributions

M.A. initiated the study. M.A, R.K., N.M., and S.S. designed experiments. M.A., R.K., A.S., and T.K. performed food modification experiment. M.A., Yo.N., and D.M. performed scRNA-seq experiment. M.A. and Ya.N. performed library preparation with advice from N.O. M.A. analyzed all data. M.A. and S.S. wrote the manuscript.

## Disclosure

The authors declare that they have no conflict of interest associated with this manuscript.

## Figure legends

**Fig. S1.**
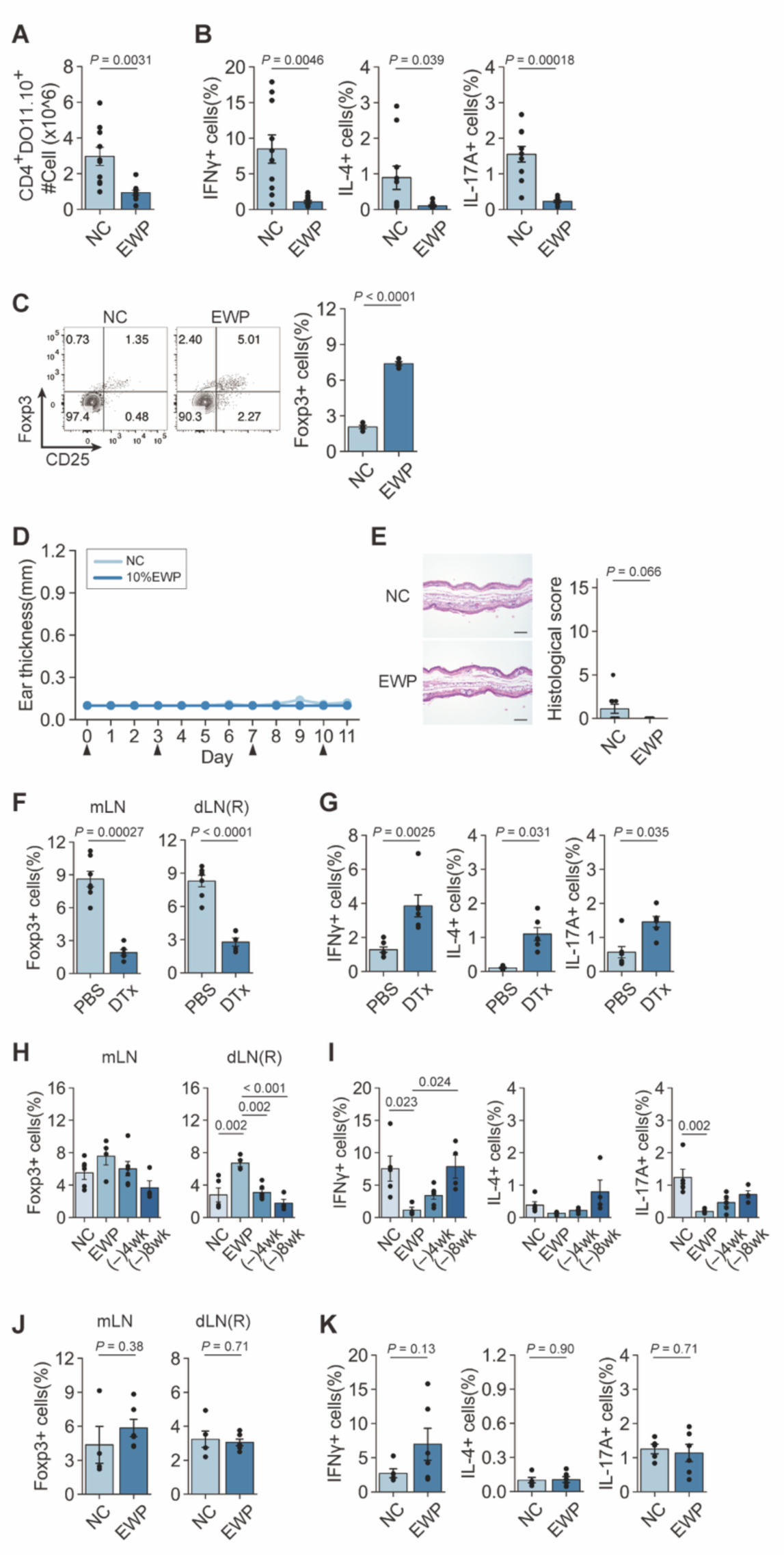
pTreg generation and maintenance by antigen feeding. (**A**) The number of DO CD4^+^ T cells in dLNs in Fig. 1A. (**B**) Frequency of IFN-γ^+^, IL-4^+^, and IL-17A^+^ cells among DO CD4^+^ T cells in dLNs (*n* = 10 and 11). (**C**) CD25 and Foxp3 expression by DO CD4^+^ T cells in the spleen at 4 weeks after designated feeding (*n* = 4). (**D**) Changes in left ear thickness in Fig. 1A. Arrowheads indicate OVA challenges. (**E**) Histology and histological scores of the left ear in NC- and EWP-fed DORe mice. Bars in histology indicate 100 µm. (**F**, **G**) Frequencies of pTreg cells in mLNs, dLNs, and the right ear (F) and frequencies of IFN-γ^+^, IL-4^+^, and IL-17A^+^ cells in dLNs (G) in the mice shown in Fig. 1G (*n* = 7 (PBS-treated) or *n* = 6 (DTx-treated)). (**H**, **I**) Frequencies of pTreg cells in mLNs and dLNs (H), and frequencies of IFN-γ^+^, IL-4^+^, and IL-17A^+^ cells in dLNs (I) in the mice shown in Fig. 1J (*n* = 4-6). (**J**, **K**) Frequencies of pTreg cells in mLNs and dLNs (J), and frequencies of IFN-γ^+^, IL-4^+^, and IL-17A^+^ cells in dLNs (K) (*n* = 4-6). Vertical bars indicate mean ± SEM. Statistical significance was assessed by unpaired *t* test or Welch’s *t* test (A, B, C, E, F, G, J, K), and Tukey-Kramer test (H, I).

**Fig. S2.**
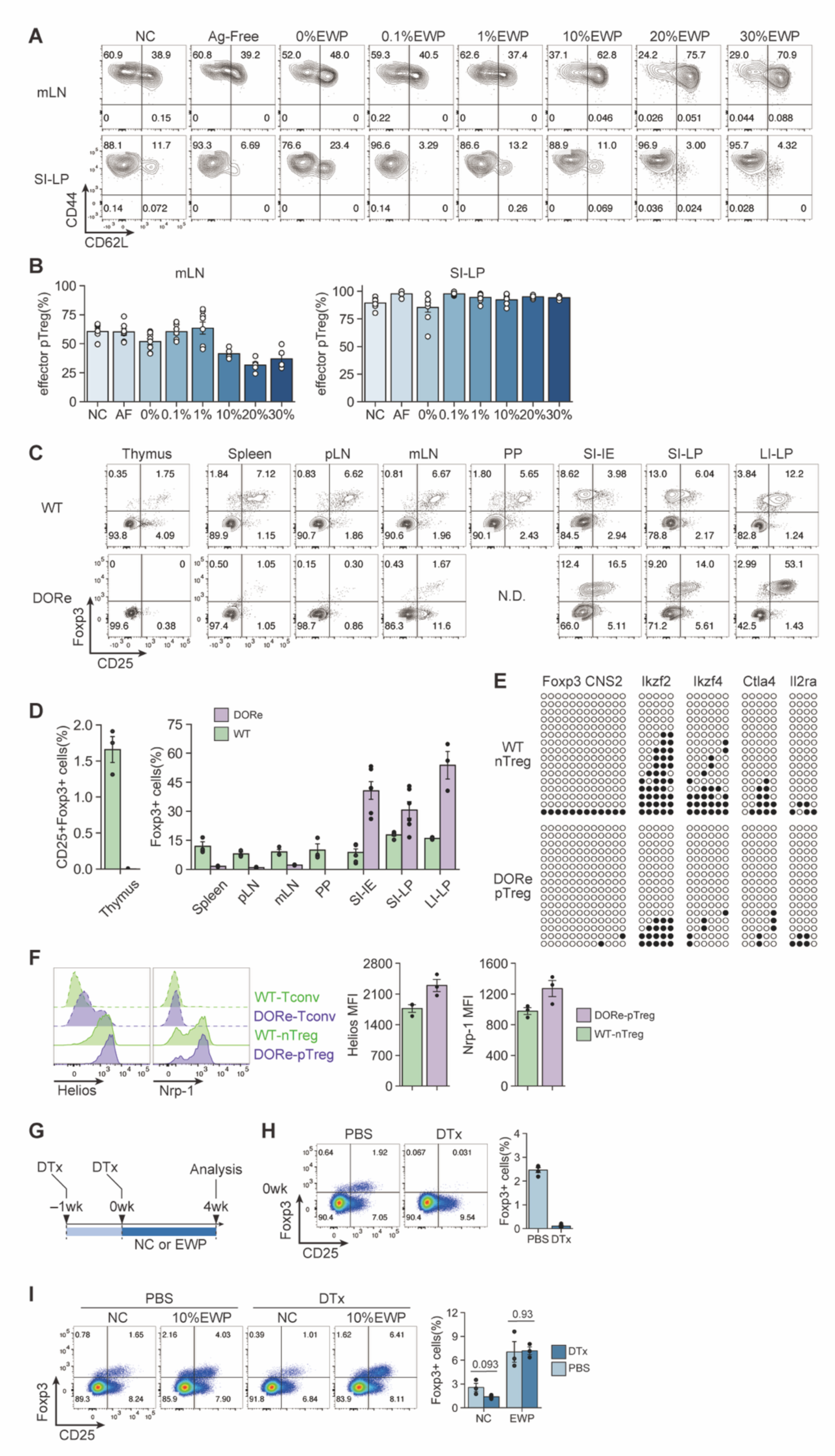
pTreg cell development at different antigen doses and after depletion of pre-existing pTreg cells in NC-fed DORe mice. (**A**, **B**) CD62L and CD44 expression by pTreg cells (A) and frequencies of CD44^hi^CD62L^lo^ effector pTreg cells (B) in mLNs and SI-LP from DORe mice fed with designated diets for 6 weeks. (**C, D**) Foxp3^+^CD4^+^ T cells in the thymus and peripheral lymphoid tissues of NC-fed DORe or WT (eFox) mice. Representative staining for Foxp3/CD25 and the frequencies of CD25^+^Foxp3^+^ T cells in the thymus and Foxp3^+^ T cells in various tissues (*n* = 3 for thymus, spleen, pLNs, mLNs, and Peyer’s patches (PP)) or (*n* = 6 for SI-IE, SI-LP, and LI-LP). **(E**) Methylation status in TSDRs of DORe-derived pTreg cells compared to eFox-derived nTreg cells. Empty and black circles indicate demethylated and methylated CpG residues, respectively. (**F**) Helios and Nrp-1 expression levels of eFox-Tconv, DORe-Tconv, eFox-nTreg, and DORe-pTreg cells. Barplots indicate MFI of Helios and Nrp-1 expression on eFox-nTreg cells and DORe-pTreg cells (*n* = 3). (**G**) *De novo* pTreg development after depletion of pTreg cells preexisting in NC-fed mice. DORF mice having been maintained with NC feeding were DTx-treated and then fed with NC or EWP-chow for 4 weeks. (**H, I**) Foxp3^+^CD4^+^ T cells immediately after DTx treatment (H) and after EWP or NC feeding for 4 weeks (I). Representative flow cytometry plots and the ratios of Foxp3^+^ pTreg cells in mLNs (*n* = 3-4). Vertical bars indicate mean ± SEM. Statistical significance was assessed by unpaired *t* test (I).

**Fig. S3.**
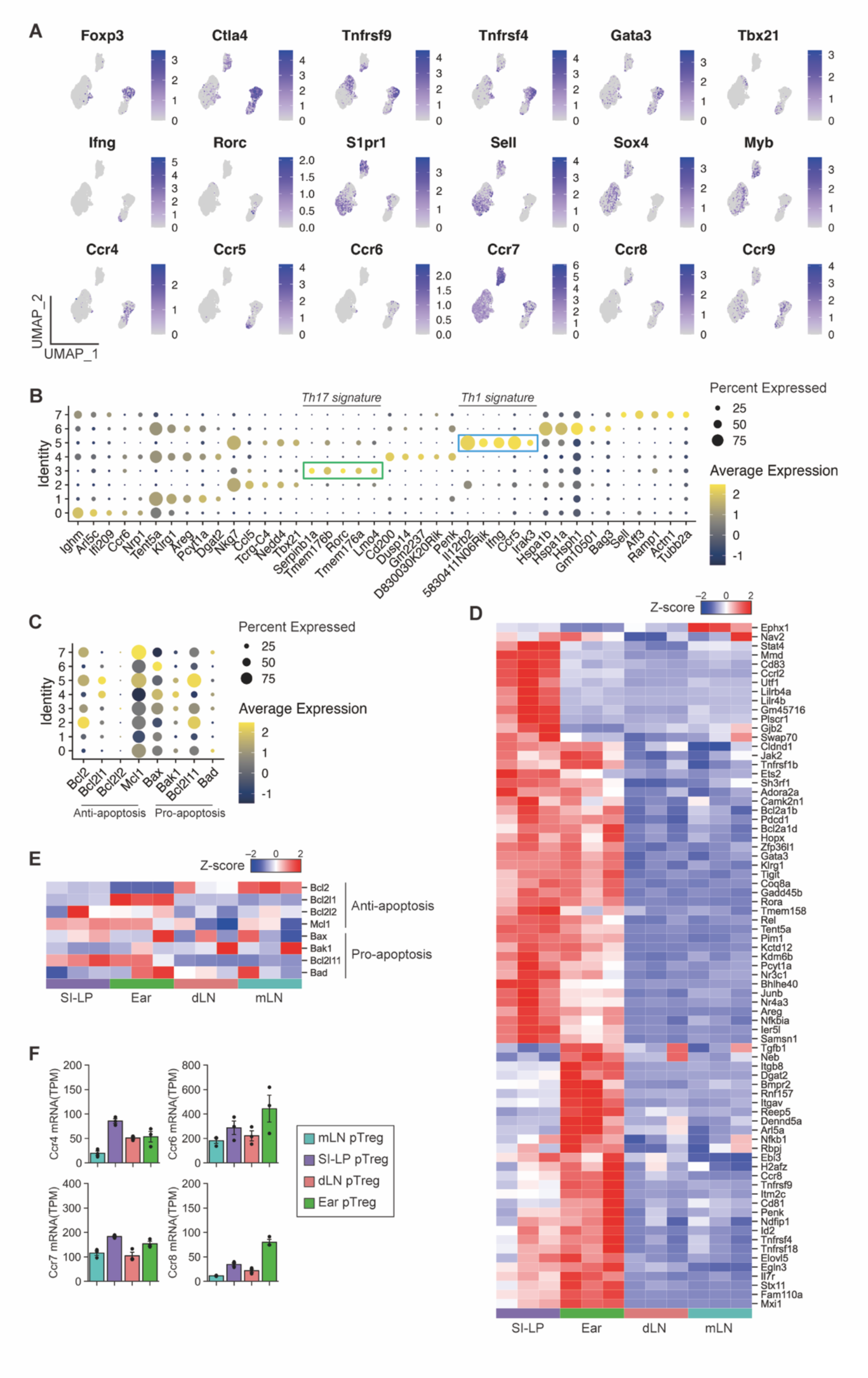
Single-cell RNA-seq analysis of T cells in DORe mice and bulk mRNA-seq analysis of pTreg cells from different tissues after EWP feeding. (**A**) Uniform manifold approximation and projection (UMAP) plots for expression levels of T cell-related molecules in DO CD4^+^ T cells in NC- or EWP-fed DORe mice. (**B**) Marker dotplots of differentially expressed genes in re-clustered T cell clusters in Fig. 3E. (**C**) Marker dotplots of anti-apoptosis and pro-apoptosis related genes in re-clustered T cell clusters. (**D**) Projection of 75 Cluster 3-up-regulated genes detected by scRNA-seq as shown in Fig. 3C on the transcriptome. (**E**) Heatmap of anti-apoptosis and pro-apoptosis related genes among pTreg cells from designated tissues. (**F**) TPM-normalized gene expression of chemokine receptors among DORe-pTreg cells derived from different tissues. Vertical bars indicate mean ± SEM.

**Fig. S4.**
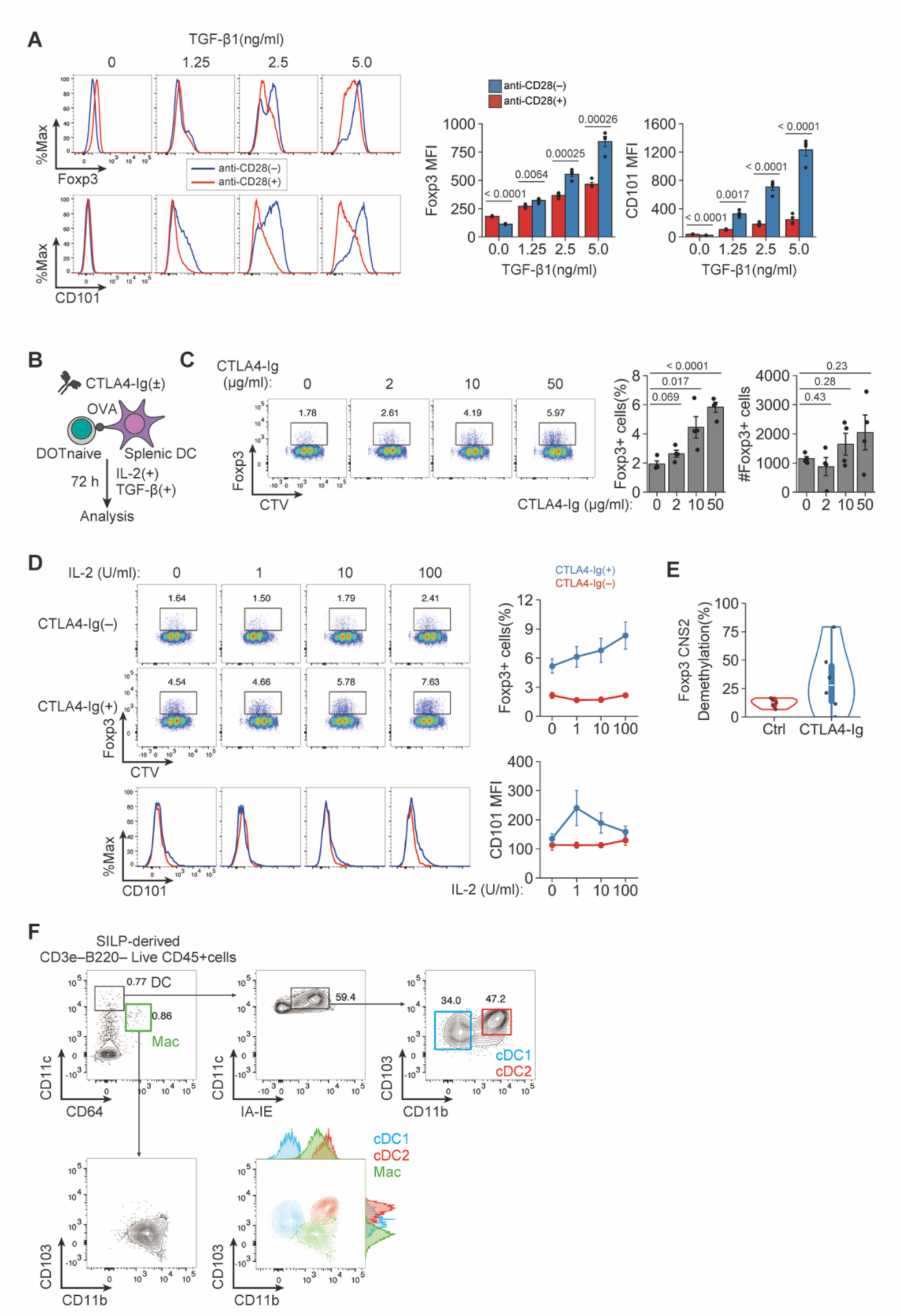
Dose effects of CTLA-4-Ig, TGF-β, and IL-2 on iTreg generation. (**A**) Synergistic effect of TGF-β supplementation and CD28 signal deprivation on *in vitro* iTreg induction and CD101 expression by naïve CD4^+^ Tconv cells (*n* = 4). (**B**) Experimental design. DORe-derived naïve Tconv cells and splenic DCs were co-cultured with or without CTLA4-Ig in the presence of OVA, TGF-β and IL-2. (**C**) Response to various concentrations of CTLA4-Ig (0, 2, 10, and 50 µg/ml) in *in vitro* iTreg induction (*n* = 4). (**D**) Response to various concentrations of IL-2 (0, 1, 10, and 100 U/ml) in *in vitro* iTreg induction and CD101 expression (*n* = 4 (0 U/ml) and *n* = 7 (1-100 U/ml)). (**E**) Degrees of demethylation as percentage at *Foxp3*-CNS2 regions of iTreg cells induced with or without CTLA4-Ig. (**F**) Gating strategy for isolating classic dendritic cells from SI-LP of WT mice. Vertical bars indicate mean ± SEM. Statistical significance was assessed by unpaired *t* test or Welch’s *t* test (A, C).

**Fig. S5.**
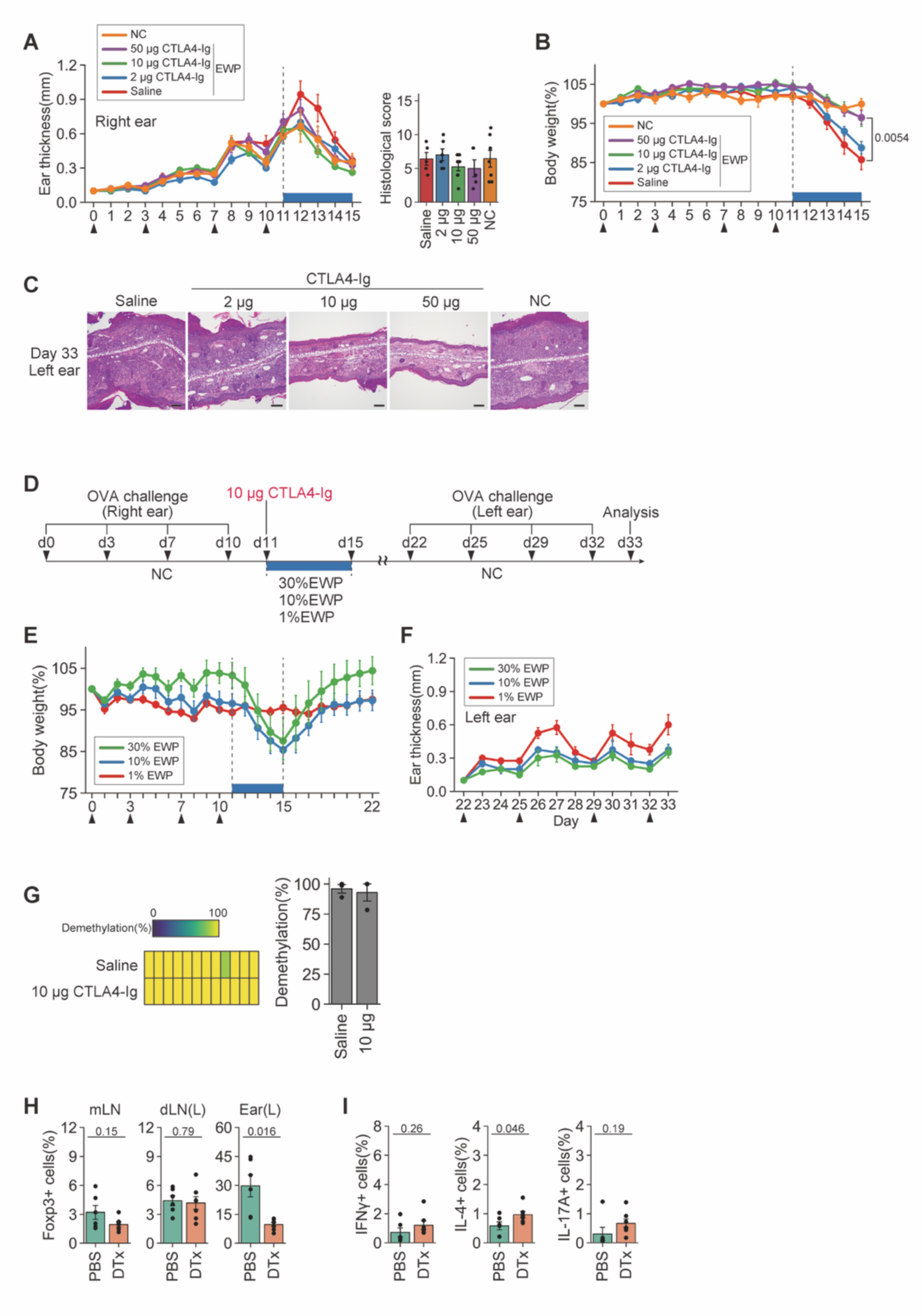
Effects of CD80/CD86 blockade and subsequent antigen feeding on antigen-presensitized mice. (**A**, **B**) Changes in ear thickness of the right ear (A) and body weight (B) of the mice treated as shown in Fig. 7A. Arrowheads indicate OVA challenges. (**C**) Representative histologies of the left ear of the mouse groups at day 33 as shown in Fig. 7C. Bars in histology indicate 100 µm. (**D-F**) Experimental protocol to assess the effects of EWP doses on oral tolerance induction by CTLA4-Ig treatment and EWP feeding (D). DORe mice were OVA sensitized at the right ear four times, i.p. injected with CTLA4-Ig on day 11, fed with 1, 10, or 30% EWP chow for four days, OVA challenged at the left ear four times from day 22, and assessed for body weight (E) and ear swelling (F) (*n* = 4). (**G**) Demethylation status and frequencies of demethylation at *Foxp3*-CNS2 locus of DORe pTreg cells isolated from the mice treated with saline or 10 µg CTLA4-Ig followed by EWP feeding and assessed at day 15 in Fig. 7A. (**H, I**) Frequencies of pTreg cells in mLNs, dLNs, and left ears (H) and of IFN-γ^+^, IL-4^+^, and IL-17A^+^ cells in dLNs at day 33 in DORF mice treated as described in Fig. 7J (I) (*n* = 6-7). Vertical bars indicate mean ± SEM. Statistical significance was assessed by unpaired *t* test or Welch’s *t* test (H, I).

**Table S1.**
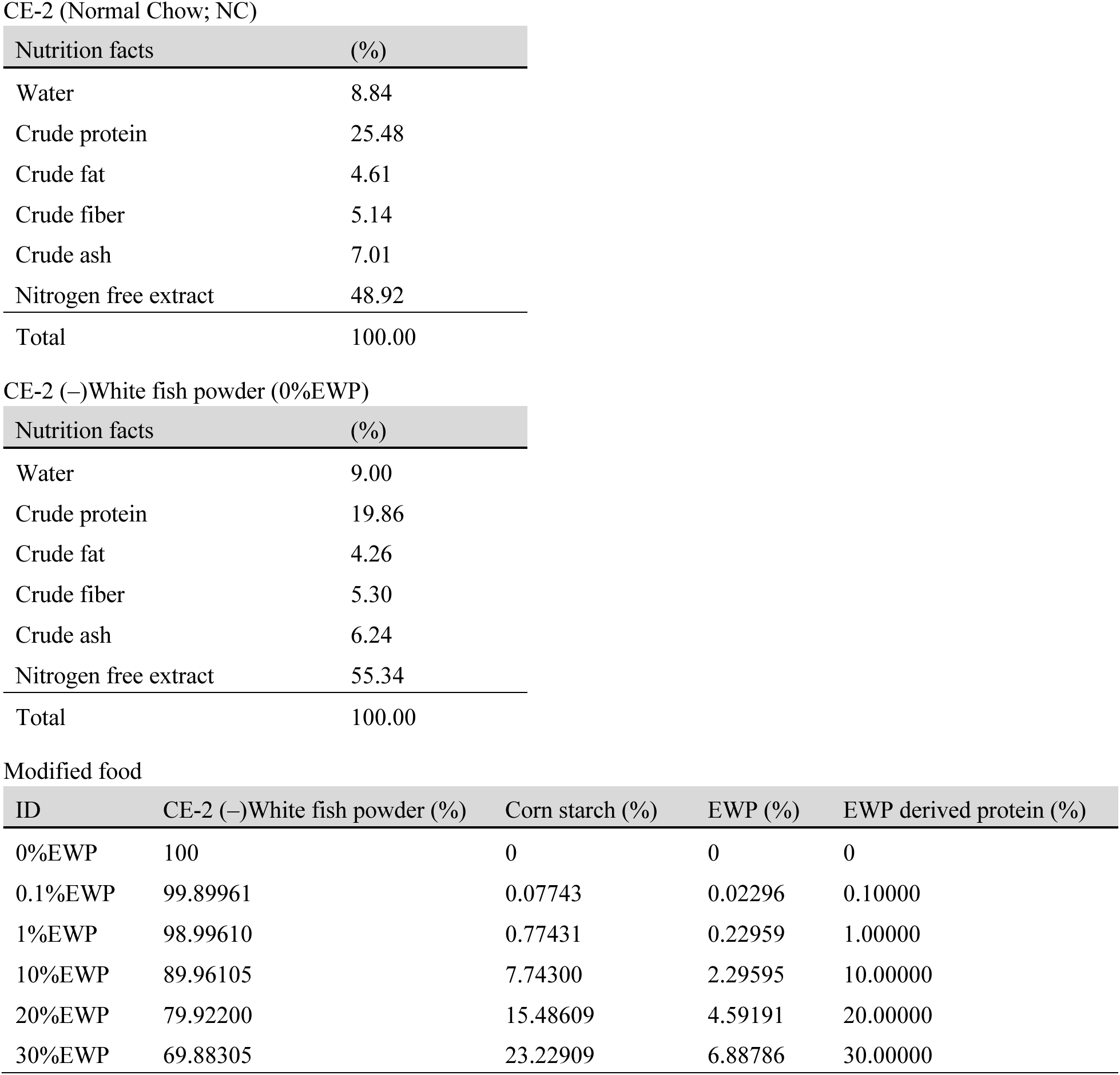
Nutrition facts of modified food.

CE-2 (Normal Chow; NC)
| Nutrition facts | (%) |
| --- | --- |
| Water | 8.84 |
| Crude protein | 25.48 |
| Crude fat | 4.61 |
| Crude fiber | 5.14 |
| Crude ash | 7.01 |
| Nitrogen free extract | 48.92 |
| Total | 100.00 |

CE-2 (-)White fish powder (0%EWP)
| Nutrition facts | (%) |
| --- | --- |
| Water | 9.00 |
| Crude protein | 19.86 |
| Crude fat | 4.26 |
| Crude fiber | 5.30 |
| Crude ash | 6.24 |
| Nitrogen free extract | 55.34 |
| Total | 100.00 |

Modified food
| ID | CE-2 (-)White fish powder (%) | Corn starch (%) | EWP (%) | EWP derived protein (%) |
| --- | --- | --- | --- | --- |
| 0%EWP | 100 | 0 | 0 | 0 |
| 0.1%EWP | 99.89961 | 0.07743 | 0.02296 | 0.10000 |
| 1%EWP | 98.99610 | 0.77431 | 0.22959 | 1.00000 |
| 10%EWP | 89.96105 | 7.74300 | 2.29595 | 10.00000 |
| 20%EWP | 79.92200 | 15.48609 | 4.59191 | 20.00000 |
| 30%EWP | 69.88305 | 23.22909 | 6.88786 | 30.00000 |

**Table S2.** List of antibodies.

| Antibodies | Manufacturer | Identifier |
| --- | --- | --- |
| Anti-mouse CD3e-APC (145-2C11) | BD pharmingen | Cat#553066; RRID: AB_398529 |
| Anti-mouse CD3e-APC-Cy7 (145-2C11) | BioLegend | Cat#100330; RRID: AB_1877170 |
| Anti-mouse CD3e-Biotin (145-2C11) | BD Biosciences | Cat#553060; RRID: AB_394593 |
| Anti-mouse CD3e-PE (145-2C11) | Invitrogen | Cat#12-0031-82; RRID: AB_465496 |
| Anti-mouse CD3e-V500 (500A2) | BD Biosciences | Cat#560771; RRID: AB_1937314 |
| Anti-mouse CD4-AF700 (RM4-5) | Invitrogen | Cat#56-0042-82; RRID: AB_494000 |
| Anti-mouse CD4-V500 (RM4-5) | BD Biosciences | Cat#560782; RRID: AB_1937327 |
| Anti-mouse CD4-BV650 (RM4-5) | BD Biosciences | Cat#563747; RRID: AB_2716859 |
| Anti-mouse CD8a-PerCP-Cy5.5 (53-6.7) | BD Biosciences | Cat#551162; RRID: AB_394081 |
| Anti-mouse CD11b-AF700 (M1/70) | Invitrogen | Cat#56-0112-80; RRID: AB_657585 |
| Anti-mouse CD11b-APC (M1/70) | BioLegend | Cat#101212; RRID: AB_312795 |
| Anti-mouse CD11b-PerCP-Cy5.5 (M1/70) | Invitrogen | Cat#45-0112-82; RRID: AB_468714 |
| Anti-mouse CD11c-BV605 (N418) | BioLegend | Cat#117334; RRID: AB_2562415 |
| Anti-mouse CD11c-PE-Cy7 (N418) | BioLegend | Cat#117318; RRID: AB_493568 |
| Purified anti-mouse CD16/32 (93) | BioLegend | Cat#101302; RRID: AB_312801 |
| Anti-mouse CD25-BV421 (PC61) | BioLegend | Cat#102034; RRID: AB_11203373 |
| Anti-mouse CD25-PE (PC61) | BD pharmingen | Cat#553866; RRID: AB_395101 |
| Anti-mouse CD25-PE-Cy7 (PC61) | BD Biosciences | Cat#552880; RRID: AB_394509 |
| Anti-mouse CD25-PerCP-Cy5.5 (PC61) | BD Biosciences | Cat#551071; RRID: AB_394031 |
| Anti-mouse CD44-APC (IM7) | Invitrogen | Cat#17-0441-83; RRID: AB_469390 |
| Anti-mouse CD44-PE-Cy7 (IM7) | Invitrogen | Cat#25-0441-82; RRID: AB_469623 |
| Anti-mouse CD44-V500 (IM7) | BD Biosciences | Cat#560780; RRID: AB_1937328 |
| Anti-mouse CD45-BV510 (30-F11) | BioLegend | Cat#103138; RRID: AB_2563061 |
| Anti-mouse CD45.1-APC-Cy7 (A20) | BioLegend | Cat#110716; RRID: AB_313505 |
| Anti-mouse/human CD45R/B220-APC-eFluor780 (RA3-6B2) | Invitrogen | Cat#47-0452-82; RRID: AB_1518810 |
| Anti-mouse/human CD45R/B220-Biotin (RA3-6B2) | BioLegend | Cat#103204; RRID: AB_312989 |
| Anti-mouse CD62L-BV421 (MEL-14) | BD Biosciences | Cat#562910; RRID: AB_2737885 |
| Anti-mouse CD62L-PerCP-Cy5.5 (MEL-14) | BD pharmingen | Cat#560513; RRID: AB_10611578 |
| Anti-mouse CD64-PE (X54-5/7.1) | BioLegend | Cat#139304; RRID: AB_10612740 |
| Anti-mouse CD90.1-BV605 (OX-7) | BD Biosciences | Cat#740374; RRID: AB_2740106 |
| Anti-mouse CD90.1-PerCP (OX-7) | BD Biosciences | Cat#557266; RRID: AB_396611 |
| Anti-mouse CD90.2-AF700 (53-2.1) | BioLegend | Cat#140324; RRID: AB_2566740 |
| Anti-mouse CD90.2-PE-Cy7 (53-2.1) | BioLegend | Cat#140310; RRID: AB_10643586 |
| Anti-mouse CD101-APC (REA301) | Miltenyi Biotec | Cat#130-104-304; RRID: AB_2654344 |
| Anti-mouse CD101-PE (REA301) | Miltenyi Biotec | Cat#130-120-173; RRID: AB_2752032 |
| Anti-mouse CD101-PE-Vio770 (REA301) | Miltenyi Biotec | Cat#130-104-305; RRID: AB_2654345 |
| Anti-mouse CD103-APC (2E7) | Invitrogen | Cat#17-1031-82; RRID: AB_1106992 |
| Anti-mouse CD103-PE-Cy7 (2E7) | BioLegend | Cat#121426; RRID: AB_2563691 |
| Anti-mouse CD134(OX40)-BV421 (OX-86) | BioLegend | Cat#119411; RRID: AB_10962569 |
| Anti-mouse CD134(OX40)-PE (OX-86) | BioLegend | Cat#119410; RRID: AB_2207344 |
| Anti-mouse CD137(4-1BB)-APC (17B5) | BioLegend | Cat#106110; RRID: AB_2564297 |
| Anti-mouse CD195(CCR5)-BV421 (C34-3448) | BD Biosciences | Cat#743695; RRID: AB_2741677 |
| Anti-mouse CD199(CCR9)-PE (CW-1.2) | BioLegend | Cat#128710; RRID: AB_1227479 |
| Anti-mouse CD278(ICOS)-PE-Cy7 (C398.4A) | BioLegend | Cat#313520; RRID: AB_10643411 |
| Anti-mouse CD304(Neupilin-1)-eFluor450 (3DS304M) | Invitrogen | Cat#48-3041-82; RRID: AB_2574051 |
| Anti-mouse CD304(Neupilin-1)-PE-Cy7 (3DS304M) | Invitrogen | Cat#25-3041-82; RRID: AB_2573436 |
| Anti-mouse CD357(GITR)-PE-Cy7 (DTA-1) | BD Biosciences | Cat#558140; RRID: AB_647252 |
| Anti-mouse/human Helios-APC (22F6) | BioLegend | Cat#137218; RRID: AB_10660750 |
| Anti-mouse/human Helios-PE-Cy7 (22F6) | BioLegend | Cat#137236; RRID: AB_2565990 |
| Anti-mouse I-A/I-E-BV421 (M5/114.15.2) | BioLegend | Cat#107631; RRID: AB_10900075 |
| Anti-mouse I-A/I-E-BV510 (M5/114.15.2) | BioLegend | Cat#107635; RRID: AB_2561397 |
| Anti-mouse/human Ki-67-PE-Cy7 (SolA15) | Invitrogen | Cat#25-5698-82; RRID: AB_11220070 |
| Anti-mouse/human KLRG1-PerCP-Cy5.5 (2F1/KLRG1) | BioLegend | Cat#138418; RRID: AB_2563014 |
| Anti-mouse/human T-bet-BV421 (4B10) | BioLegend | Cat#644816; RRID: AB_2686976 |
| Anti-mouse TCR DO11.10-APC (KJ1-26) | Invitrogen | Cat#17-5808-80; RRID: AB_469459 |
| Anti-mouse TCR DO11.10-PerCP-Cy5.5 (KJ1-26) | BioLegend | Cat#118512; RRID: AB_2028522 |
| Anti-mouse TER-119/Erythroid Cells-Biotin (TER-119) | BD pharmingen | Cat#553672; RRID: AB_394985 |
| Anti-mouse TIGIT-APC (1G9) | BioLegend | Cat#142105; RRID: AB_10960139 |
| Anti-mouse Foxp3-Alexa Fluor 488 (FJK-16s) | Invitrogen | Cat#53-5773-82; RRID: AB_763537 |
| Anti-human/mouse Gata3-eFluor660 (TWAJ) | Invitrogen | Cat#50-9966-42; RRID: AB_10596663 |
| Anti-mouse RORgamma(t)-PE (B2D) | Invitrogen | Cat#12-6981-82; RRID: AB_10807092 |
| Anti-mouse IL-4-APC (11B11) | Invitrogen | Cat#17-7041-82; RRID: AB_469494 |
| Anti-mouse IL-17A-BV421 (TC11-18H10.1) | BioLegend | Cat#506925; RRID: AB_10900442 |
| Anti-mouse IFN gamma-PE (XMG1.2) | Invitrogen | Cat#12-7311-82; RRID: AB_466193 |
| TotalSeq-C0301 anti-mouse Hashtag 1 Antibody | BioLegend | Cat#155861; RRID: AB_2800693 |
| TotalSeq-C0302 anti-mouse Hashtag 2 Antibody | BioLegend | Cat#155863; RRID: AB_2800694 |
| TotalSeq-C0303 anti-mouse Hashtag 3 Antibody | BioLegend | Cat#155865; RRID: AB_2800695 |
| TotalSeq-C0304 anti-mouse Hashtag 4 Antibody | BioLegend | Cat#155867; RRID: AB_2800696 |
| Purified NA/LE anti-mouse CD3e (145-2C11) | BD pharmingen | Cat#553057; RRID: AB_394590 |
| Purified NA/LE anti-mouse CD28 (37.51) | BD pharmingen | Cat#553294; RRID: AB_394763 |
| Anti-Histone H3 Acetyl Lys27 | GeneTex | Cat#GTX60815; RRID: AB_2888004 |

## Notes

### Competing Interest Statement

The authors have declared no competing interest.

### Summary of Updates

Added new Figure 5 and changed the format.

